# Loss of Corneal Raver2 Expression in Aniridia Associated Keratopathy

**DOI:** 10.1101/2020.02.21.959585

**Authors:** Subrata K. Das, Lara Carroll, Derick Holt, Xiaohui Zhang, Hironori Uehara, Bonnie Archer, Jayakrishna Ambati, Balamurali K. Ambati

**Affiliations:** School of Medical Science and Technology, IIT Kharagpur-721302, India; Moran Eye Center, University of Utah, Salt Lake City, Utah- 84132, USA; Loma Linda University Medical Center, Loma Linda, CA-92350, USA; Center for Advanced Vision Science, University of Virginia, Charlottesville, VA- 22908 USA

**Keywords:** Raver2, Pax6, aniridia, aniridia-associated keratopathy

## Abstract

**Purpose:** The aim of this study was to evaluate downstream factors of Pax6 associated with the aniridic corneal phenotype. We characterize the ocular expression of *Raver2*, a novel heterogeneous ribonucleoprotein (hnRNP).

**Methods:** Genome wide microarrays of corneal RNA from non-aniridic mice (C57BL/6J and MRL/MpJ) and corneas from aniridia model, Pax6^+/-^ (SEY) mice were performed. *Raver2* expression profiles were validated using qPCR and western blot. Raver2 immunohistochemistry on corneas of Pax6^+/-^ mice and aniridic humans were compared to control specimens, and insitu-hybridization was used to analyze developmental expression of *Raver2*.

**Results:** Microarray analysis of corneal RNA identified *Raver2* as the gene with the strongest differential expression profile across non-aniridic and aniridic mouse strains. Consistent with mouse data, we found that *Raver2* is expressed at high levels in healthy human corneal epithelium, and is downregulated in the corneas of human aniridia patients.

**Conclusions:** These findings suggest that *Raver2* has an evolutionarily conserved developmental role downstream of *Pax6* in corneal development and maintenance.

## Introduction

A clear, transparent cornea is essential for vision. Loss of transparency in the cornea is the second leading cause of global blindness and is caused by a variety of disorders such as ocular hypoxia, inflammation, corneal trauma, alkali injury, and limbal stem cell deficiency^1-5^. Congenital aniridia is a rare ocular disorder that may affect as many as 1 in 64,000 to 100,000 people worldwide^6, 7^ with defects arising in the cornea, anterior chamber, iris, lens, retina, macula, and the optic nerve. Approximately 20% of aniridia patients develop aniridia-associated keratopathy (AAK)^8-12^. In this progressive pathology, signs of keratopathy appear early in life, with thickening and vascularization of the peripheral cornea which may gradually advance to pathologic corneal neovascularization (KNV) of the entire cornea^10^. The majority of aniridia cases result from a varied assortment of autosomal dominant loss-of function mutations in the Paired box 6 gene (*Pax6*) encoding a phylogenetically conserved transcription factor with central roles in the organization of the developing eye as well as the post-natal maintenance of a transparent, avascular corneal epithelium^13-18^. The Pax6^+/-^ (SEY) mouse carries a heterozygous mutation deriving from a spontaneous G to T transversion in codon 194 of the *Pax6* locus, and recapitulates key features of human aniridia. This transversion replaces glycine with a stop codon, resulting in peptide termination prior to the homeobox domain and subsequent *Pax6* haploinsufficiency, spontaneous KNV, and a spectrum of aniridia related pathologies with a progression in corneal disease severity that parallels AAK in humans^19^.

In contrast to the corneal pathologies found in Pax6^+/-^ mice, the MRL/MpJ mouse strain, known as the “super-healing” mouse, responds to corneal alkali burn (a corneal wounding model) with rapid reepithelialization, reduced inflammation, fibrotic remodeling, and no loss of corneal transparency^20^. Intermediate to these two strains, the C57BL/6 mouse is susceptible to corneal insult, responding to alkali wounding with robust inflammatory cell infiltration, extensive scarring, ulceration, neovascularization and loss of transparency caused by scar tissue ^21^. We explored the differential expression profiles of these mice and found that Raver2 is strongly downregulated in corneal epithelium of Pax6^+/-^ mutants relative to C57Bl6 and MRL/MpJ strains. *Raver2* is a novel hnRNP protein with three N-terminal RNA Recognition Motif (RRM) domains. Although no known biological function has been reported for *Raver2*, it was identified as a homologue of another hnRNP, *Raver1*, based on domain architecture, sequence homology, and similarity within N-terminal RRM domains^22^. Like *Raver1, Raver2* associates with its well characterized RNA-binding co-factor, polypyrimidine tract binding protein (PTB) and is thought to stabilize PTB assembly on endogenous RNAs enabling recruitment of the protein/RNA complexes from the cytoplasm to the perinucleolar compartment^23^. Several hnRNPs function in mRNA stabilization, processing and splicing^24^. Herein we present our findings correlating loss of *Raver2* expression in the corneal epithelium with Pax6-associated defects.

## Methods

### Animals

Male and female C57BL/6J (stock no. 000664), MRL/MpJ (stock no. 000486), and B6EiC3Sn a/A-Pax6Sey /J (Pax6+/-, stock no. 000391) mice were purchased from The Jackson Laboratory (Bar Harbor, ME). Experimental groups were age and sex matched. All the mice were handled in accordance with the Association for Research in Vision and Ophthalmology (ARVO) Statement for the use of Animals in Ophthalmic and Vision Research. Experiments were approved by the Institutional Animal Care and Use Committees (IACUCs) of the University of Utah.

### Corneal Microarray and Data Analysis

Corneas were harvested immediately after euthanizing 8-week old female C57BL/6, MRL/MpJ, and Pax6+/- mice and transferred immediately into RNAlater stabilization agent (Qiagen, Valencia, CA, USA). Corneas were trimmed of any remaining limbus or iris, and total RNA was extracted with the RNeasy Micro Kit (Qiagen). For each sample, 50ng of total RNA, plus oligo dT/T7 RNA polymerase promoter oligonucleotide primers and MMLV-RT RNA polymerase were used to synthesize cDNA before transfer to the University of Utah Microarray Core Facility. Samples were processed using an Agilent G2505B Microarray Scanner (http://www.hci.utah.edu/sharedrsrs/CoreGroup.srs?deptId=33). TIF files generate from the scan were loaded into Agilent Feature Extraction Software version 10.1.1.1, which automatically positions a grid and finds the centroid position of each feature on the array, calculates feature intensities and background, then records data as a tab-delimited text file. Each Cy3 or Cy5 hybridization was treated as an individual biological replicate in subsequent data analysis. Microarray intensity data was filtered to remove control features and any features flagged as non-uniform or feature population outliers. Log2 intensity data from all samples and all genes was clustered in R using Ward’s method and Euclidean distance. Heat-maps were generated in R using the heatmap.2 function in the gplots library from BioConductor. Genes correlated or anti-correlated with Raver2 expression were clustered by first calculating the mean expression value for each gene, and then calculating the deviation from the mean for each gene in each sample. These deviations were hierarchically clustered using Euclidean distance and complete linkage. The color scale represents deviation from mean expression for each gene, with increased expression displayed in red and decreased expression in green.

### cDNA synthesis and Quantitative RT-PCR (qPCR)

cDNAs were synthesized from total RNA (corneal) using the Omniscript RT kit (Qiagen) with oligo-dT (dT20) primers according to the manufacturer’s protocol. Real-time qPCR was performed with the QuantiTect SYBR Green PCR Kit (Qiagen) with amplification on a GeneAmp 5700 Thermocycler (ABI, Foster City, CA). Expression levels were normalized to internal control gene GAPDH using the delta delta CT method^25^.

### Immunostaining of Human Corneal Tissue

Human tissue use was approved by the respective Institutional Review Boards and conformed to the Declaration of Helsinki. Normal human donor corneal tissue was obtained from the Utah Lions Eye Bank and underwent paraffin embedding followed by sectioning and staining for Raver2. Paraffin blocks of corneal specimens known to have a pre-existing diagnosis of aniridia were obtained from the research collections of the University of Utah and the University of Kentucky and underwent sectioning and staining. For immuno histochemical staining, deparaffinized sections underwent antigen retrieval using antigen unmasking solution (Vector, H-3300). Endogenous peroxidase was quenched with 0.3% H202 prepared in methanol. Samples were then incubated with 5% donkey serum block (Sigma D9663). Immunolocalization was performed with Raver2 polyclonal antibody (1:50; Santa Cruz, sc-165338), or Raver1 polyclonal antibody (1:50; Santa Cruz, sc-102073) followed by donkey anti-goat IgG-HRP conjugated secondary antibody (Santa Cruz, sc-2020) and counterstained with hematoxylin (Sigma-Aldrich, GHS232). Each specimen was ranked ordinally by a blind observer according to signal intensity in corneal epithelium for comparison among the normal and anirida groups (1=lowest <2 <3 <4=highest).

### Cell Culture, siRNA treatment and Western Blotting

HCECs (ATCC, cat# PCS-700-010) were cultured in serum free medium (ATCC, cat# PCS-700-030) supplemented with growth factors according to the manufacturer’s instructions. To prevent loss of epithelial cell properties, cultures were limited to passages four through seven. siRNAs targeting Raver2, Pax6 and non-specific control siRNA were used (predesigned FlexiTube siRNAs, Qiagen). For siRNA transfection, 2×10^5^ cells/well (6-well plate) HCECs were transfected with 30pmol siRNA using lipofectamine RNAiMax (Life Technologies, Grand Island, NY, USA) according to the manufacturer’s protocol. After 48 hrs, cultured cell lysate were made in RIPA buffer. Samples were run on a 10% SDS-PAGE gel, and Western blotting was performed using standard techniques with Raver2 (Santa Cruz) and Pax6 (Abcam, ab195045) antibodies. The same protocol was used for cell lysate preparation and Western blotting analysis from mouse cornea.

### RNA in situ hybridization

Corneas were harvested immediately after euthanizing mice and fixed in 4% paraformaldehyde (PFA) for 2 hours at 4°C. Cryoprotected frozen sections (10µm) were permeabilized by 5µg/ml proteinase-K treatment for 5 minutes postfixed in 4% PFA for 10 minutes and acetylated with 0.1 M triethanolamine (TEA) buffer, pH 8.0, containing 0.25% (v/v) acetic anhydride (Sigma) 4% for 10 minutes. Following a 10 minute prehybridization (4× SSC containing 50% (v/v) deionized formamide at RT°), DIG-labeled Raver2 anti-sense or sense (control) RNA probes (0.5 µg per ml) were applied in hybridization buffer (40% deionized formamide. 10% dextran sulfate, 1× Denhardt’s solution, 4× SSC, 10 mM DTT, 1 mg/ml yeast t-RNA, 1 mg/ml denatured, sheared salmon sperm DNA and heat denatured RNA probe) overnight in a humidified chamber at 42°C, after which sections were washed at 37°C for 30 min in 2XSSC, for 30 min in 1X SSC, and treated with 20 µg/ml RNase A for 30 mins at 37°C to digest any single-stranded (unbound) RNA probe. After post-hybridization washes (twice in 0.2 X SSC at 37), DIG labeled probes were visualized with anti-DIG alkaline phosphatase conjugate, 5-bromo-4-chloro-3-indolylphosphate (BCIP) and nitroblue tetrazolium chloride (NBT) according to the manufacturer’s protocol (Roche Applied Science, Indianapolis, IN, USA). The sections were then mounted. Images were obtained with an EVOS FL Auto Cell Imaging System (LifeTechnology, Bothell,WA).

### Statistics

For microarrary analysis, normalized and corrected fluorescence intensity values were averaged across replicates for each microarray probe to yield a single value for each probe sequence (and genotype). Values were log2-transformed and quantile normalized. Normalized data was uploaded to GeneSifter (www.geospiza.com) for differential expression analysis. Differentially expressed genes were selected using ANOVA, requiring at least 2-fold differential expression and a Benjamini and Hochberg-corrected p value < 0.05. For qPCR and corneal imaging, differences between groups were tested using Students t-test or ANOVA. A value of P < 0.05 is considered statistically significant for all analyses.

## Results

### Raver2 expression is strongly downregulated in corneas of Pax6^+/-^ mice

Genome-wide mRNA expression in corneas of Pax6^+/-^ (*SEY*), C57Bl6, and MRL/MpJ mice was analyzed by microarray, using n=5-6 independent replicates of each strain. Seventy-six genes were strongly downregulated in Pax6 corneas relative to C57Bl6 and MRL/MpJ corneas, (Figure 1A, B and Supplemental Table 1) whereas 96 genes were strongly upregulated in Pax6^+/-^ corneas relative to C57Bl6 and MRL/MpJ corneas (Supplemental Figure 1 and Supplemental Table 2). Whole genome microarray data clustered tightly among biological replicates of the same strain (Supplemental Figure 2). We focused on the genes with low expression in Pax6^+/-^ mice and high expression in C57Bl6 and MRL/MpJ mice. The gene with the strongest differential expression profile across the three corneal models was *Raver2*, which was downregulated 9.6 and 14.4 fold in Pax6^+/-^ corneas relative to C57Bl/6J and MRL/MpJ respectively. Quantitative RT-PCR (q-PCR) and western blot validated the Raver2 microarray expression data and suggested that *Raver2* parallels *Pax6* expression across the three corneal models (Figure 1C and D). We then tested whether in vitro downregulation of *Pax6* in cultured human epithelial cells (HCEC) similarly reduced *Raver2* expression. In contrast to the strong downregulation of *Raver2* in *Pax6* haploinsufficient corneas, *Raver2* expression was maintained in HCECs following transfection with *Pax6* siRNA. Similarly, *Pax6* expression was maintained following the converse transfection (*Raver2* siRNA), suggesting an indirect regulatory association between these two factors in corneal epithelium (Supplemental Figure 3).

**Figure 1.**
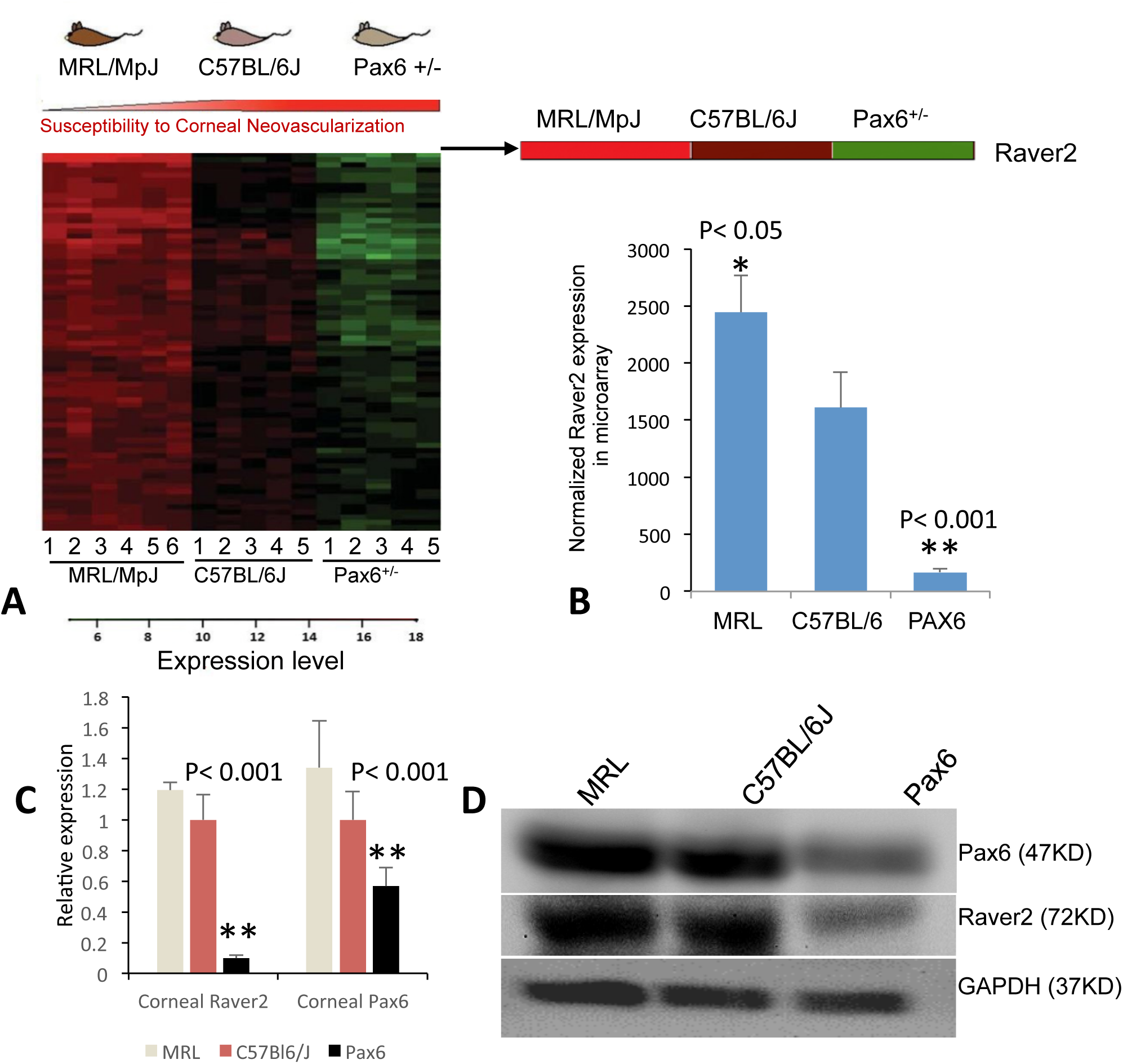
Di□erential gene expression of corneas from three mouse strains. (A) Genome-wide cDNA microarray identifies *Raver2* as the top di□erentially expressed gene in the corneas of three mouse strains. Seventy six genes were identified with high, intermediate, and low expression in MRL/MpJ, C57BL/6J, and Pax6+/- respectively. Log (2)-transformed expression data for all biological replicates is expressed as a heat map for all genes within this group. Each row represents a specific oligonucleotide probe on the array and each column represents an independent biological replicate with strain indicated below the heat map. (B) Normalized absolute intensities show downregulation of Raver2 expression in Pax6 corneas (9.6-fold) and upregulation in MRL/MpJ (1.5-fold) relative to C57BL/6J. (C, D) Downregulation of Raver2 and Pax6 expression in Pax6 corneas by (C) qPCR and (D) Western blot.

**Figure 2.**
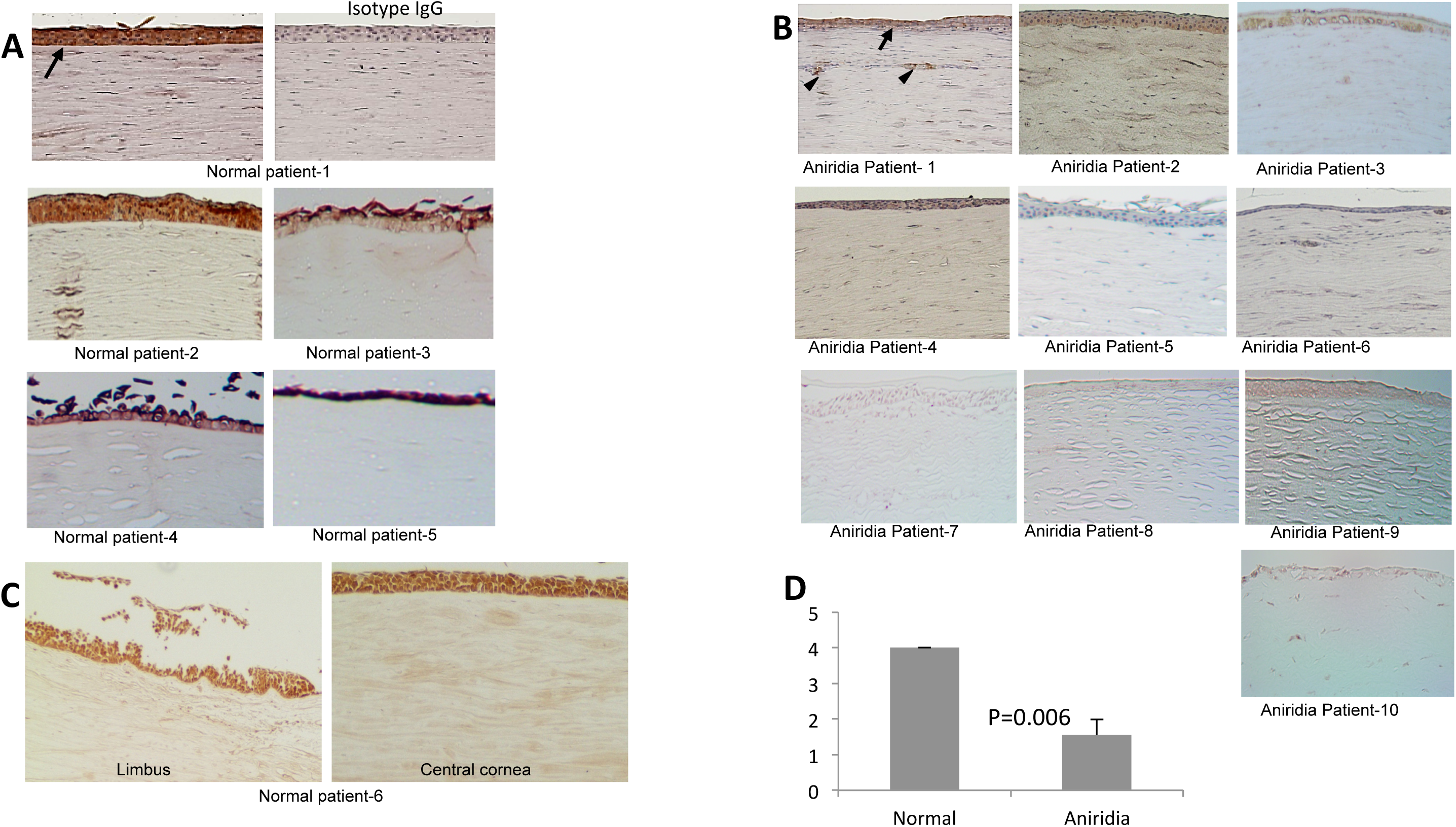
*Raver2* is expressed at high levels in normal human corneal epithelium and diminished in patients with aniridia. (A) Raver2-specific antibody demonstrates strong staining within normal human corneal epithelium, whereas no signal is seen using isotype control antibody. (B) Human aniridia corneal specimens removed at the time of corneal transplantation. Raver2 specific staining is significantly reduced within corneal epithelium. Specimens show hallmarks of AAK, including vascularization (black arrowheads) and lack of regular Bowman’s membrane and epithelial thinning (arrows). All slides used hematoxylin counterstain. Magnification is 10X for all photomicrographs. (C) Limbus and central cornea from the same normal patient shows similar levels of staining. (D) Quantification of brown color intensity on a scale of 1 to 4. (n = 6 normal cornea and n = 10 aniridia cornea. Values are means ± SEM).

**Figure 3.**
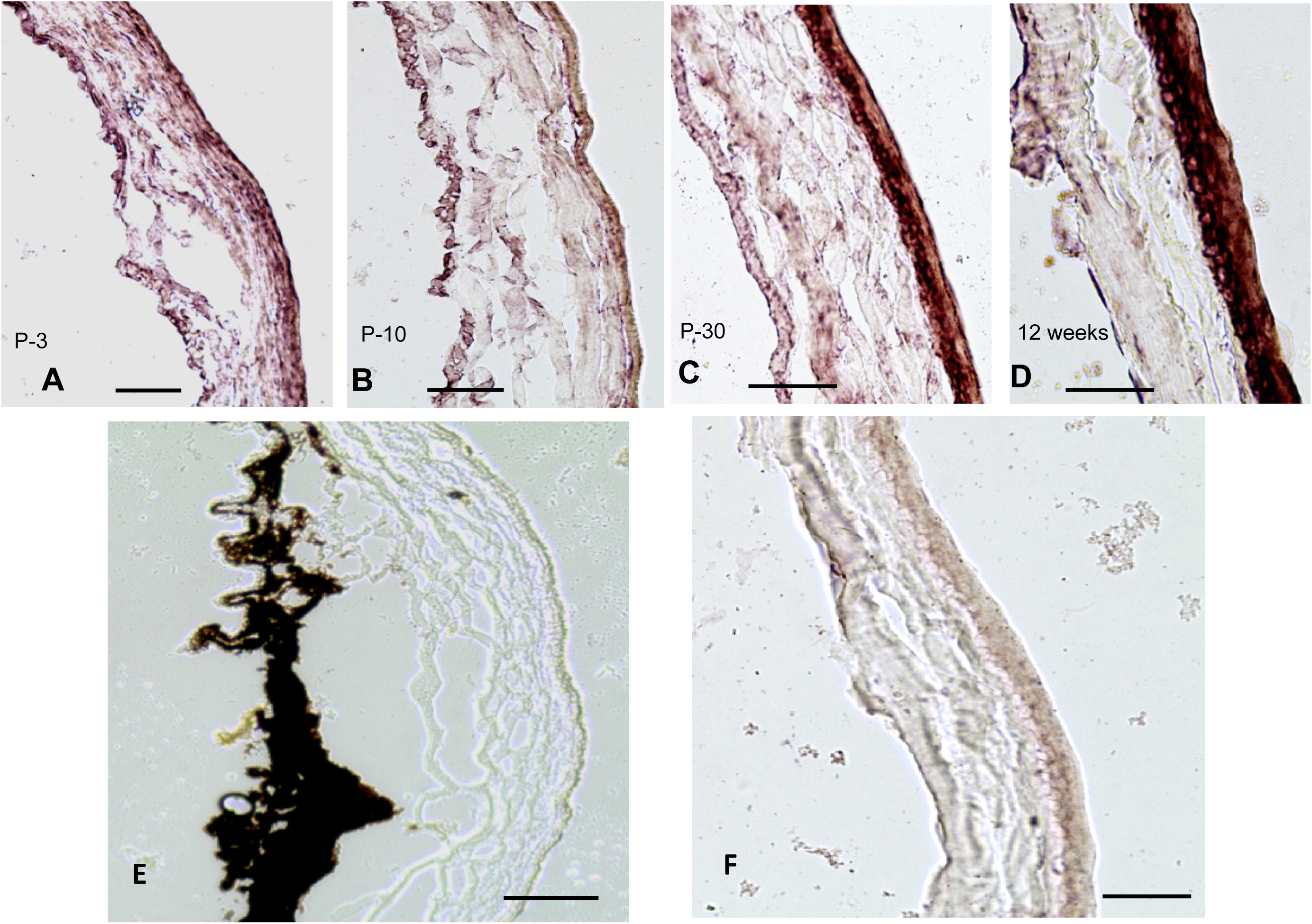
RNA *in-situ* hybridization with Raver2 anti-sense probe. (A-D) C57BL/6J corneas at different developmental time points showing gradual upregulation of Raver2 in epithelium from P-3 to adult using a Raver2 anti-sense probe. (E) Raver2 anti-sense probe does not label adult Pax6 mouse cornea. (F) Adult C57BL/6J cornea labeled with sense probe. Scale bar = 50µm.

### Raver2 expression is downregulated in aniridic corneas

Human corneal specimens from normal donors and patients with aniridia were analyzed using immunohistochemistry to determine Raver2 expression levels. Normal corneas showed strong Raver2 expression localized primarily within corneal epithelium extending throughout central and limbal regions, but it is also expressed at low levels in corneal stroma. Raver2 signal in aniridic corneas was much less prominent (Figure 2, and Supplemental Table 3). RNA *In-Situ* hybridization with a *Raver2* antisense probe revealed developmental progression of *Raver2* corneal expression in mice, showing a gradual upregulation as the cornea matures from birth to P30 (Figure 3). *Raver2* expression is maintained at high levels in C57Bl/6J adult corneal tissue, but is absent in the corneas of Pax6^+/-^ animals. These data collectively suggest that diminished *Raver2* expression is associated with aniridic corneal pathology. To determine if *Raver1* expression might compensate for *Raver2* loss in Pax6 haploinsufficient corneas, we examined Raver1 protein in corneas of aniridic humans and mice using immunohistochemistry. Localization of this hnRNP family member overlapped with that of Raver2 in corneal epithelium, however, levels of Raver1 expression were unchanged between healthy vs aniridic corneas in humans, and between C57Bl6 and Pax6 haploinsufficient mice (Figure 4), suggesting that *Raver1* does not compensate for *Raver2* deficiency.

**Figure 4.**
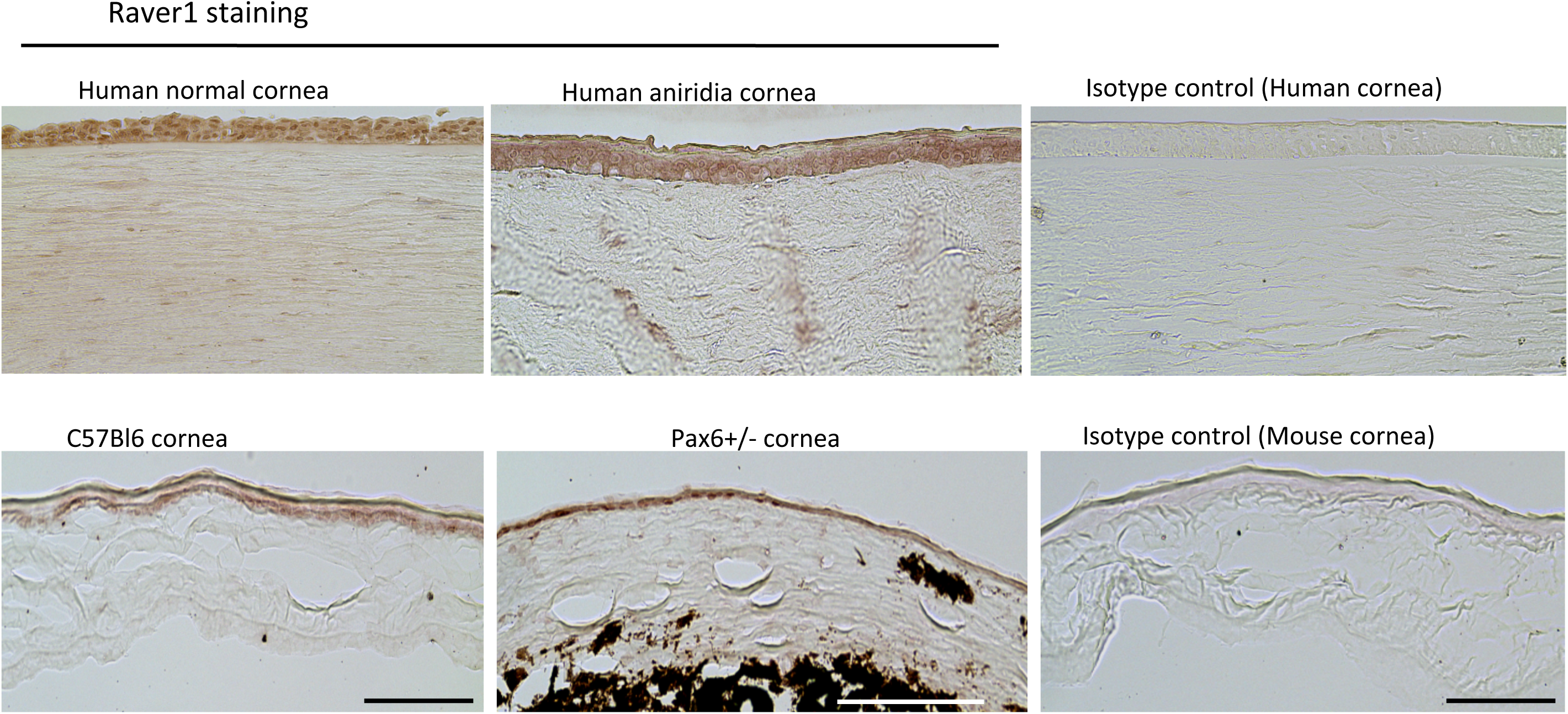
Raver1 is expressed in human and mouse cornea. Representative images of normal human cornea and aniridia cornea showing Raver1 localization in corneal epithelium. Raver1 is also expressed in normal C57Bl/6 and Pax6 mouse cornea. Scale bar =100µm.

## Discussion

We used genome wide microarray to identify novel genes that function as regulators of corneal homeostasis. Using mice with and without Pax6 haploinsufficiency, we identified Raver2 as the top differentially expressed gene candidate. Pax6^+/-^ mice share many features of aniridia with human patients that carry a defective Pax6 gene. The corneas of both aniridia patients and Pax6^+/-^ mice were found to express low levels of *Raver2* in contrast to the high levels of *Raver2* expression found in healthy corneas from humans, C57Bl/6 and MRL/MpJ mice. As *Raver2* corneal expression increases from birth to adulthood, these combined data suggest that Raver2 has an evolutionarily conserved developmental role in the cornea, and that Raver2 deficiency may contribute to the pathogenesis of vision-threatening AAK.

Like *Pax6*^26^, corneal *Raver2* expression is highly expressed in epithelium where it is specifically downregulated in both Pax6 haploinsufficient mice and aniridia patients. Interestingly, isolated cultures of healthy HCECs did not exhibit *Raver2* downregulation after transfection with *Pax6* specific siRNA, nor did we conversely find downregulation of *Pax6* expression after *Raver2* specific siRNA transfection. This result strongly suggests an indirect regulatory relationship between *Pax6* and *Raver2*. Pax6 is a multifunctional protein regulating differentiation via expression of different downstream molecules in a highly context dependent manner^27-32^. Normal eye development and maintenance depends on Pax6 dosage, with Pax6 haploinsufficiency and overexpression leading to overlapping sets of anterior and posterior segment defects^33-35^. Coordination of the timing and dose of *Pax6* expression is critical for proper differentiation and development of the ectodermally derived cornea, which may confound efforts to replicate molecular events in epithelial cell culture. Alternatively, regulatory cross-talk between *Pax6* and *Raver2* may require signaling input from other corneal cell types, such as the corneal stroma, or infiltrating leukocytes that respond to corneal damage, which were not present in our cell culture. Similarly to Raver1, Raver2 interacts with its well characterized co-factor, PTB, suggesting it functions in post-transcriptional regulation^36^. Mechanistically, the function of Raver2 may be to recruit and/or stabilize PTB assembly on mRNA of key angiogenic genes, working in concert to repress or promote splicing, intron retention and early polyadenylation, as has previously been described for *Raver1* in the regulation of alpha-tropomyosin splicing^37-39^. Intron retention can also regulate the expression of cellular genes by modulating mRNA stability, subcellular localization, and/or translation, which are known functions performed by PTB in cooperation with other hnRNPs^40, 41^. Moreover, recent evidence that PTB can bind directly to U1 snRNA ^42^ suggests that distinct RNA processing pathways can directly interact to influence spliceosomal assembly and ultimately determine mRNA isoform selection. Future studies will focus on delineating the molecular mechanisms of *Pax6* regulation of *Raver2*, limbal stem cell formation, and corneal morphogenesis. A worthy area of investigation would be to explore whether therapeutic intervention upregulating *Raver2* can prevent the progression or development of such pathologies.

## Acknowledgments

We acknowledge J.A. and B.K.A. who are the co-founder of iVeena Holdings, iVeena Delivery Systems and Inflammasome Therapeutics, and also J.A. has been a consultant for Allergan, Biogen, Boehringer-Ingelheim, Immunovant, Janssen, Olix Pharmaceuticals, Retinal Solutions, and Saksin LifeSciences unrelated to this work. We would like to thank Deepak Edward, Anthony Aldave, Clark Springs and Ralph C. Eagle Jr. for providing corneas of Aniridia patients. This work was funded by NEI 5R01EY17950 (B.K.A).

## Figure Legends

Figure S1. Microarray expression profiles of 97 genes upregulated in Pax6+/- and downregulated in MRL/MpJ. Log(2)-transformed expression data for all biological replicates is expressed as a heat map for all genes within this group. Each row represents a specific oligonucleotide probeon the array and each column represents an independent biological replicate with strain indicated below the heat map. A complete list of genes within this group is given in Supplemental Table 1and 2.

Figure S2. Genome-wide microarray data clusters according to mouse strain. Whole microarray datasets were subjected to unbiased clustering analysis using Ward’s method. The data clustere dtightly according to mouse strain, demonstrating excellent data qualityand reproducibility. MRL/MpJ and C57BL/6J corneal expression patterns were closer to one another than either was to Pax6+/-.

Figure S3. Western blot of Pax6 and Raver2 showing Human Corneal Epithelial cells (HCEC) treated with siRNA targeting Pax6 and Raver2, negative siRNA and No siRNA as a mock control.

## References

1. Azar DT. Corneal angiogenic privilege: angiogenic and antiangiogenic factors in corneal avascularity, vasculogenesis, and wound healing (an American Ophthalmological Society thesis). Trans Am Ophthalmol Soc. 2006;104:264-302. PubMed PMID: 17471348; PubMed Central PMCID: PMC1809914.

2. Beebe DC. Maintaining transparency: a review of the developmental physiology and pathophysiology of two avascular tissues. Semin Cell Dev Biol. 2008;19(2):125–33. doi: 10.1016/j.semcdb.2007.08.014. PubMed PMID: 17920963; PubMed Central PMCID: PMC2276117.

3. Carmeliet P. Angiogenesis in life, disease and medicine. Nature. 2005;438(7070):932–6. doi: 10.1038/nature04478. PubMed PMID: 16355210.

4. Dua HS, Azuara-Blanco A. Limbal stem cells of the corneal epithelium. Surv Ophthalmol. 2000;44(5):415-25. PubMed PMID: 10734241.

5. Papas EB. The role of hypoxia in the limbal vascular response to soft contact lens wear. Eye Contact Lens. 2003;29(1 Suppl):S72-4; discussion S83-4, S192-4. PubMed PMID: 12772736.

6. Eden U, Iggman D, Riise R, Tornqvist K. Epidemiology of aniridia in Sweden and Norway. Acta Ophthalmol. 2008;86(7):727–9. doi: 10.1111/j.1755-3768.2008.01309.x. PubMed PMID: 18494745.

7. Nelson LB, Spaeth GL, Nowinski TS, Margo CE, Jackson L. Aniridia. A review. Surv Ophthalmol. 1984;28(6):621-42. PubMed PMID: 6330922.

8. Brandt JD, Casuso LA, Budenz DL. Markedly increased central corneal thickness: an unrecognized finding in congenital aniridia. Am J Ophthalmol. 2004;137(2):348–50. doi: 10.1016/j.ajo.2003.09.038. PubMed PMID: 14962429.

9. Hingorani M, Hanson I, van Heyningen V. Aniridia. Eur J Hum Genet. 2012;20(10):1011–7.doi: 10.1038/ejhg.2012.100. PubMed PMID: 22692063; PubMed Central PMCID: PMC3449076.

10. Lee H, Khan R, O’Keefe M. Aniridia: current pathology and management. Acta Ophthalmol. 2008;86(7):708–15. doi: 10.1111/j.1755-3768.2008.01427.x. PubMed PMID: 18937825.

11. Lopez-Garcia JS, Garcia-Lozano I, Rivas L, Martinez-Garchitorena J. [Congenital aniridia keratopathy treatment]. Arch Soc Esp Oftalmol. 2006;81(8):435-44. PubMed PMID: 16933167.

12. Tseng SC, Li DQ. Comparison of protein kinase C subtype expression between normal and aniridic human ocular surfaces: implications for limbal stem cell dysfunction in aniridia. Cornea. 1996;15(2):168-78. PubMed PMID: 8925665.

13. Jordan T, Hanson I, Zaletayev D, Hodgson S, Prosser J, Seawright A, Hastie N, van Heyningen V. The human PAX6 gene is mutated in two patients with aniridia. Nat Genet. 1992;1(5):328–32. doi: 10.1038/ng0892-328. PubMed PMID: 1302030.

14. Ton CC, Hirvonen H, Miwa H, Weil MM, Monaghan P, Jordan T, van Heyningen V, Hastie ND, Meijers-Heijboer H, Drechsler M, et al. Positional cloning and characterization of a paired box- and homeobox-containing gene from the aniridia region. Cell. 1991;67(6):1059-74. PubMed PMID: 1684738.

15. Yokoi T, Nishina S, Fukami M, Ogata T, Hosono K, Hotta Y, Azuma N. Genotype-phenotypem correlation of PAX6 gene mutations in aniridia. Hum Genome Var. 2016;3:15052. doi: 10.1038/hgv.2015.52. PubMed PMID: 27081561; PubMed Central PMCID: PMC4760117.

16. Gehring WJ. The master control gene for morphogenesis and evolution of the eye. Genes Cells. 1996;1(1):11-5. PubMed PMID: 9078363.

17. Halder G, Callaerts P, Gehring WJ. Induction of ectopic eyes by targeted expression of the eyeless gene in Drosophila. Science. 1995;267(5205):1788-92. PubMed PMID: 7892602.

18. Gregory-Evans CY, Wang X, Wasan KM, Zhao J, Metcalfe AL, Gregory-Evans K. Postnatal manipulation of Pax6 dosage reverses congenital tissue malformation defects. J Clin Invest. 2014;124(1):111–6. doi: 10.1172/JCI70462. PubMed PMID: 24355924; PubMed Central PMCID: PMC3871240.

19. Ramaesh T, Collinson JM, Ramaesh K, Kaufman MH, West JD, Dhillon B. Corneal abnormalities in Pax6+/-small eye mice mimic human aniridia-related keratopathy. Invest Ophthalmol Vis Sci. 2003;44(5):1871-8. PubMed PMID: 12714618.

20. Heydemann A. The super super-healing MRL mouse strain. Front Biol (Beijing). 2012;7(6):522–38. doi: 10.1007/s11515-012-1192-4. PubMed PMID: 24163690; PubMed Central PMCID: PMC3806350.

21. Saika S. Yin and yang in cytokine regulation of corneal wound healing: roles of TNF-alpha. Cornea. 2007;26(9 Suppl 1):S70–4. doi: 10.1097/ICO.0b013e31812f6d14. PubMed PMID: 17881920.

22. Kleinhenz B, Fabienke M, Swiniarski S, Wittenmayer N, Kirsch J, Jockusch BM, Arnold HH, Illenberger S. Raver2, a new member of the hnRNP family. FEBS Lett. 2005;579(20):4254–8. doi: 10.1016/j.febslet.2005.07.001. PubMed PMID: 16051233.

23. Henneberg B, Swiniarski S, Sabine B, Illenberger S. A conserved peptide motif in Raver2 mediates its interaction with the polypyrimidine tract-binding protein. Exp Cell Res. 2010;316(6):966–79. doi: 10.1016/j.yexcr.2009.11.023. PubMed PMID: 19962980.

24. Dreyfuss G, Kim VN, Kataoka N. Messenger-RNA-binding proteins and the messages they carry. Nat Rev Mol Cell Biol. 2002;3(3):195–205. doi: 10.1038/nrm760. PubMed PMID: 11994740.

25. Livak KJ, Schmittgen TD. Analysis of relative gene expression data using real-time quantitative PCR and the 2(-Delta Delta C(T)) Method. Methods. 2001;25(4):402–8. doi: 10.1006/meth.2001.1262. PubMed PMID: 11846609.

26. Koroma BM, Yang JM, Sundin OH. The Pax-6 homeobox gene is expressed throughout the corneal and conjunctival epithelia. Invest Ophthalmol Vis Sci. 1997;38(1):108-20. PubMed PMID: 9008636.

27. Glaser T, Lane J, Housman D. A mouse model of the aniridia-Wilms tumor deletion syndrome. Science. 1990;250(4982):823-7. PubMed PMID: 2173141.

28. Hill RE, Favor J, Hogan BL, Ton CC, Saunders GF, Hanson IM, Prosser J, Jordan T, Hastie ND, van Heyningen V. Mouse small eye results from mutations in a paired-like homeobox-containing gene. Nature 1991;354(6354):522–5. doi: 10.1038/354522a0. PubMed PMID: 1684639.

29. Osumi N, Shinohara H, Numayama-Tsuruta K, Maekawa M. Concise review: Pax6 transcription factor contributes to both embryonic and adult neurogenesis as a multifunctional regulator. Stem Cells. 2008;26(7):1663–72. doi: 10.1634/stemcells.2007-0884. PubMed PMID: 18467663.

30. Sansom SN, Griffiths DS, Faedo A, Kleinjan DJ, Ruan Y, Smith J, van Heyningen V, Rubenstein JL, Livesey FJ. The level of the transcription factor Pax6 is essential for controlling the balance between neural stem cell self-renewal and neurogenesis. PLoS Genet. 2009;5(6):e1000511. doi: 10.1371/journal.pgen.1000511. PubMed PMID: 19521500; PubMed Central PMCID: PMC2686252.

31. Schedl A, Ross A, Lee M, Engelkamp D, Rashbass P, van Heyningen V, Hastie ND. Influence of PAX6 gene dosage on development: overexpression causes severe eye abnormalities. Cell. 1996;86(1):71-82. PubMed PMID: 8689689.

32. van Raamsdonk CD, Tilghman SM. Dosage requirement and allelic expression of PAX6 during lens placode formation. Development. 2000;127(24):5439-48. PubMed PMID: 11076764.

33. Collinson JM, Chanas SA, Hill RE, West JD. Corneal development, limbal stem cell function, and corneal epithelial cell migration in the Pax6(+/-) mouse. Invest Ophthalmol Vis Sci. 2004;45(4):1101-8. PubMed PMID: 15037575.

34. Collinson JM, Quinn JC, Hill RE, West JD. The roles of Pax6 in the cornea, retina, and olfactory epithelium of the developing mouse embryo. Dev Biol. 2003;255(2):303-12. PubMed PMID: 12648492.

35. Davis J, Piatigorsky J. Overexpression of Pax6 in mouse cornea directly alters corneal epithelial cells: changes in immune function, vascularization, and differentiation. Invest Ophthalmol Vis Sci. 2011;52(7):4158–68. doi: 10.1167/iovs.10-6726. PubMed PMID: 21447684; PubMed Central PMCID: PMC3175973.

36. Spellman R, Rideau A, Matlin A, Gooding C, Robinson F, McGlincy N, Grellscheid SN, Southby J, Wollerton M, Smith CW. Regulation of alternative splicing by PTB and associated factors. Biochem Soc Trans. 2005;33(Pt 3):457–60. doi: 10.1042/BST0330457. PubMed PMID: 15916540.

37. Gromak N, Rideau A, Southby J, Scadden AD, Gooding C, Huttelmaier S, Singer RH, Smith CW. The PTB interacting protein raver1 regulates alpha-tropomyosin alternative splicing. EMBO J. 2003;22(23):6356–64. doi: 10.1093/emboj/cdg609. PubMed PMID: 14633994; PubMed Central PMCID: PMC291850.

38. Joshi A, Coelho MB, Kotik-Kogan O, Simpson PJ, Matthews SJ, Smith CW, Curry S. Crystallographic analysis of polypyrimidine tract-binding protein-Raver1 interactions involved in regulation of alternative splicing. Structure. 2011;19(12):1816–25. doi: 10.1016/j.str.2011.09.020. PubMed PMID: 22153504; PubMed Central PMCID: PMC3420021.

39. Rideau AP, Gooding C, Simpson PJ, Monie TP, Lorenz M, Huttelmaier S, Singer RH, Matthews S, Curry S, Smith CW. A peptide motif in Raver1 mediates splicing repression by interaction with the PTB RRM2 domain. Nat Struct Mol Biol. 2006;13(9):839–48. doi: 10.1038/nsmb1137. PubMed PMID: 16936729.

40. Sharma S, Maris C, Allain FH, Black DL. U1 snRNA directly interacts with polypyrimidine tract-binding protein during splicing repression. Mol Cell. 2011;41(5):579–88. doi: 10.1016/j.molcel.2011.02.012. PubMed PMID: 21362553; PubMed Central PMCID: PMC3931528.

41. Yap K, Lim ZQ, Khandelia P, Friedman B, Makeyev EV. Coordinated regulation of neuronal mRNA steady-state levels through developmentally controlled intron retention. Genes Dev. 2012;26(11):1209–23. doi: 10.1101/gad.188037.112. PubMed PMID: 22661231; PubMed Central PMCID: PMC3371409.

42. Peterson ML. Mechanisms controlling production of membrane and secreted immunoglobulin during B cell development. Immunol Res. 2007;37(1):33-46. PubMed PMID: 17496345.

