## Supplementary figures and images for "Loss of Corneal Raver2 Expression in Aniridia Associated Keratopathy"

### Supplementary Figure-1

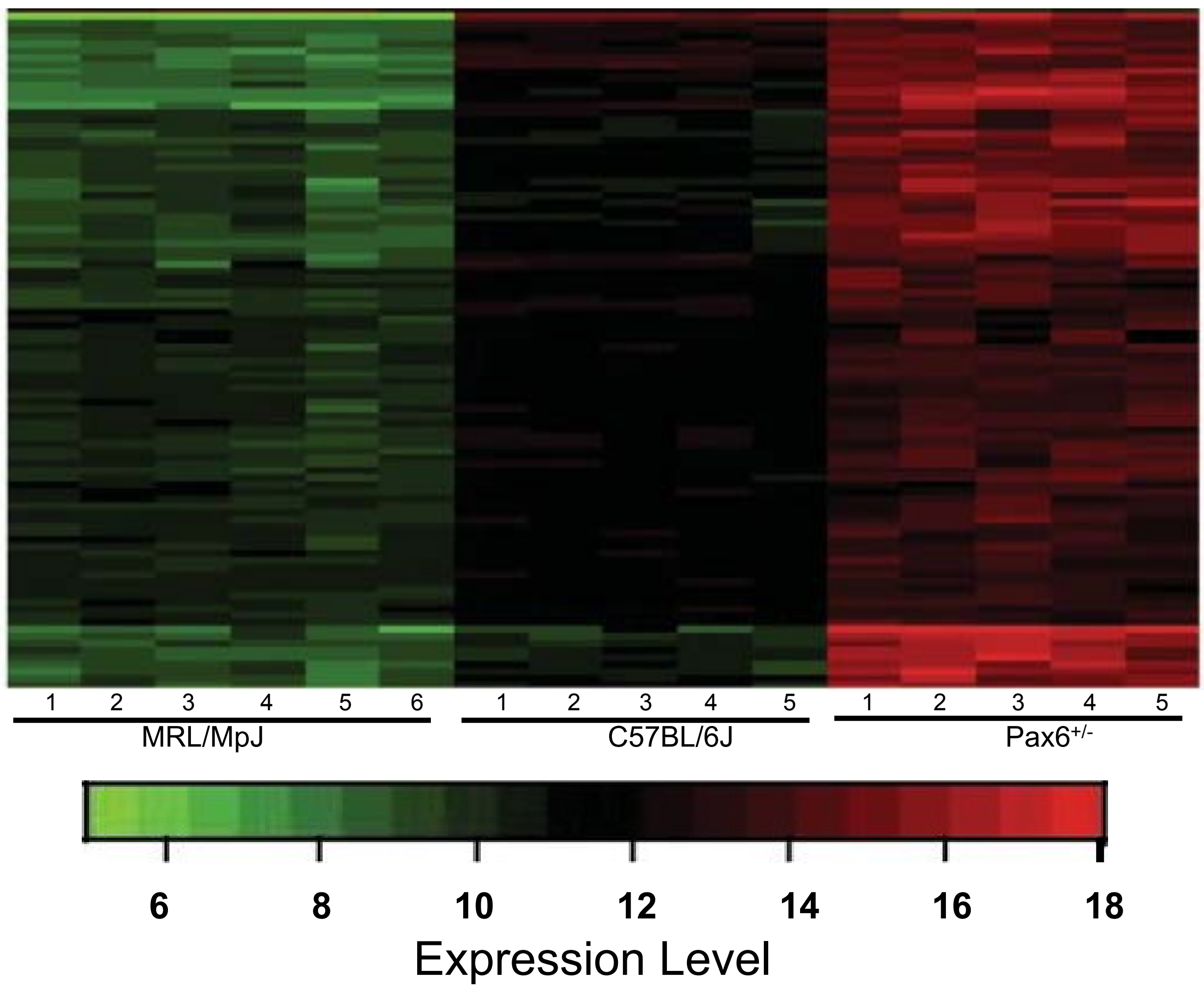

### Supplementary Figure-2

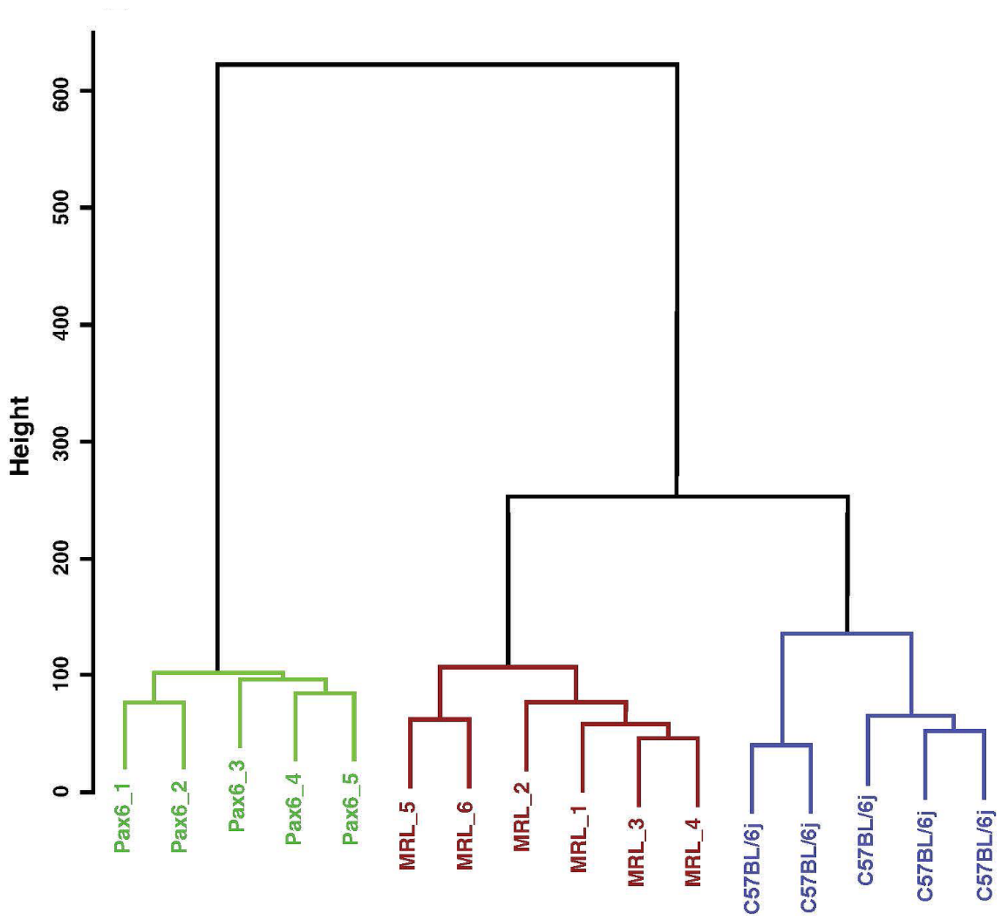

### Supplementary Figure-3

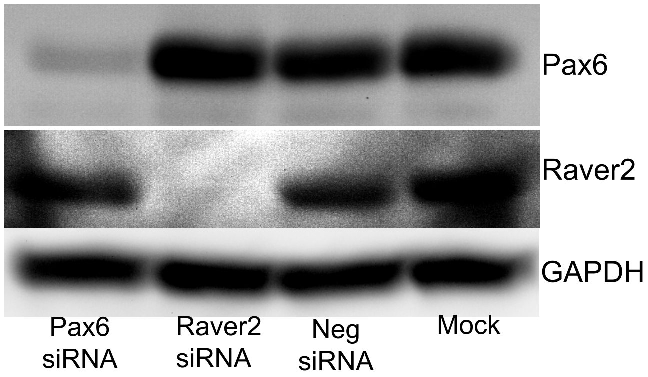
