## Supplementary Table-1 & 2 for "Loss of Corneal Raver2 Expression in Aniridia Associated Keratopathy"

Supplemental Table S1

| Group | Genbank ID | GeneName | Mean MRL log2 intensity | Mean WT log2 intensity | Mean PAX6 log2 intensity | MRL_1 log2 intensity | MRL_2 log2 intensity | MRL_3 log2 intensity | MRL_4 log2 intensity | MRL_5 log2 intensity | MRL_6 log2 intensity | WT_1 log2 intensity | WT_2 log2 intensity | WT_3 log2 intensity | WT_4 log2 intensity | WT_5 log2 intensity | PAX6_1 log2 intensity | PAX6_2 log2 intensity | PAX6_3 log2 intensity | PAX6_4 log2 intensity | PAX6_5 log2 intensity |
|---|---|---|---|---|---|---|---|---|---|---|---|---|---|---|---|---|---|---|---|---|---|
| MRL-Up | NM_176963 | Galm | 9.425 | 8.858 | 7.741 | 9.406 | 9.532 | 9.715 | 9.500 | 9.230 | 9.166 | 8.854 | 9.217 | 8.735 | 9.000 | 8.485 | 7.710 | 7.485 | 7.662 | 7.953 | 7.895 |
| MRL-Up | U21674 | Tcte3 | 8.518 | 7.963 | 6.112 | 8.438 | 8.769 | 8.661 | 8.388 | 8.345 | 8.506 | 7.728 | 8.011 | 7.966 | 8.234 | 7.877 | 6.199 | 5.998 | 5.959 | 6.122 | 6.282 |
| MRL-Up | NM_029946 | 4931407K02Rik | 12.959 | 12.429 | 10.323 | 12.894 | 13.188 | 12.950 | 12.866 | 12.742 | 13.111 | 12.431 | 12.649 | 12.305 | 12.467 | 12.292 | 10.359 | 10.548 | 9.860 | 10.227 | 10.619 |
| MRL-Up | NM_028166 | 1600014C10Rik | 14.044 | 13.515 | 12.555 | 14.042 | 14.136 | 14.112 | 13.899 | 13.964 | 14.114 | 13.511 | 13.685 | 13.331 | 13.539 | 13.509 | 12.265 | 12.535 | 12.295 | 12.816 | 12.862 |
| MRL-Up | NM_025329 | 0610012D17Rik | 12.613 | 12.002 | 10.872 | 12.542 | 12.653 | 12.515 | 12.582 | 12.594 | 12.790 | 11.764 | 11.975 | 12.254 | 11.933 | 12.085 | 11.348 | 10.723 | 10.688 | 10.468 | 11.132 |
| MRL-Up | NM_016785 | Tpmt | 9.345 | 8.688 | 7.487 | 9.310 | 9.403 | 9.410 | 9.357 | 9.277 | 9.314 | 8.522 | 8.778 | 8.624 | 8.983 | 8.530 | 7.708 | 7.232 | 7.449 | 7.306 | 7.739 |
| MRL-Up | NM_026592 | B230118H07Rik | 13.831 | 13.173 | 12.551 | 13.691 | 13.805 | 13.740 | 13.822 | 13.796 | 14.129 | 13.237 | 13.202 | 13.263 | 12.880 | 13.283 | 12.105 | 12.238 | 12.825 | 12.695 | 12.896 |
| MRL-Up | NM_170597 | Creg2 | 8.675 | 7.053 | 6.368 | 8.586 | 8.429 | 8.850 | 8.808 | 8.835 | 8.544 | 6.953 | 6.968 | 7.183 | 6.969 | 7.192 | 6.426 | 5.994 | 6.555 | 6.417 | 6.449 |
| MRL-Up | NM_008815 | Etv4 | 8.678 | 7.909 | 7.363 | 8.665 | 9.063 | 8.628 | 8.377 | 8.567 | 8.767 | 8.084 | 7.996 | 7.639 | 7.940 | 7.885 | 7.647 | 7.237 | 7.526 | 7.235 | 7.172 |
| MRL-Up | NM_026304 | l7Rn6 | 11.736 | 11.142 | 10.307 | 11.867 | 11.802 | 11.913 | 11.753 | 11.616 | 11.461 | 11.043 | 11.101 | 11.149 | 11.351 | 11.068 | 10.538 | 9.971 | 9.948 | 10.502 | 10.576 |
| MRL-Up | NM_007904 | Ednrb | 10.242 | 9.658 | 8.066 | 10.773 | 10.325 | 10.555 | 10.049 | 9.779 | 9.972 | 9.882 | 9.642 | 9.858 | 9.419 | 9.487 | 7.843 | 7.976 | 7.869 | 8.218 | 8.423 |
| MRL-Up | NM_008838 | Pigf | 12.660 | 11.787 | 10.867 | 12.638 | 12.863 | 12.573 | 12.598 | 12.569 | 12.720 | 11.679 | 11.866 | 11.786 | 11.722 | 11.884 | 11.293 | 10.707 | 10.989 | 10.593 | 10.753 |
| MRL-Up | NM_027678 | Zranb3 | 10.283 | 9.696 | 8.928 | 10.283 | 10.412 | 10.220 | 10.088 | 10.188 | 10.507 | 9.649 | 9.878 | 9.706 | 9.746 | 9.502 | 9.216 | 9.211 | 8.889 | 8.915 | 8.411 |
| MRL-Up | NM_027495 | Tmem144 | 8.060 | 7.442 | 6.584 | 8.273 | 8.195 | 8.222 | 8.019 | 7.915 | 7.738 | 7.379 | 7.631 | 7.330 | 7.698 | 7.173 | 6.748 | 6.359 | 6.688 | 6.558 | 6.569 |
| MRL-Up | NM_026470 | Spata6 | 7.633 | 6.978 | 6.466 | 7.786 | 7.733 | 7.764 | 7.500 | 7.502 | 7.515 | 6.952 | 7.109 | 6.948 | 7.227 | 6.653 | 6.248 | 6.329 | 6.689 | 6.697 | 6.369 |
| MRL-Up | NM_030021 | D730039F16Rik | 9.182 | 7.705 | 6.430 | 9.303 | 9.182 | 9.266 | 9.067 | 9.037 | 9.235 | 7.585 | 7.756 | 7.693 | 7.709 | 7.782 | 6.521 | 6.238 | 6.566 | 6.326 | 6.501 |
| MRL-Up | NM_013564 | Insl3 | 11.370 | 8.936 | 8.250 | 11.419 | 11.415 | 11.614 | 11.312 | 11.228 | 11.230 | 8.941 | 8.972 | 8.735 | 9.240 | 8.790 | 8.438 | 7.992 | 8.268 | 8.388 | 8.165 |
| MRL-Up | NM_001081155 | Rap1gap | 12.950 | 12.367 | 11.748 | 12.994 | 12.857 | 13.016 | 13.046 | 12.889 | 12.900 | 12.388 | 12.186 | 12.466 | 12.296 | 12.501 | 11.644 | 12.025 | 11.742 | 11.642 | 11.686 |
| MRL-Up | NM_021272 | Fabp7 | 8.036 | 7.193 | 6.401 | 7.716 | 7.652 | 8.155 | 8.281 | 8.449 | 7.963 | 7.097 | 7.049 | 7.145 | 7.070 | 7.604 | 6.283 | 6.261 | 6.519 | 6.279 | 6.661 |
| MRL-Up | NM_027289 | Nt5dc2 | 8.788 | 8.298 | 7.721 | 8.771 | 8.745 | 9.070 | 8.823 | 8.635 | 8.682 | 8.324 | 8.458 | 8.174 | 8.276 | 8.259 | 7.487 | 7.458 | 7.961 | 7.739 | 7.962 |
| MRL-Up | NM_026102 | Daam1 | 10.514 | 9.963 | 9.403 | 10.282 | 10.604 | 10.615 | 10.472 | 10.577 | 10.533 | 9.894 | 10.182 | 9.728 | 10.159 | 9.854 | 9.391 | 9.549 | 9.140 | 9.243 | 9.691 |
| MRL-Up | NM_020013 | Fgf21 | 7.804 | 7.285 | 5.711 | 7.726 | 8.190 | 7.802 | 7.750 | 7.577 | 7.782 | 7.212 | 7.487 | 7.294 | 7.533 | 6.897 | 5.771 | 5.543 | 6.044 | 5.600 | 5.595 |
| MRL-Up | NM_172574 | Pqlc3 | 11.778 | 10.716 | 10.021 | 11.738 | 11.969 | 11.886 | 11.789 | 11.535 | 11.754 | 10.423 | 10.815 | 10.857 | 10.904 | 10.580 | 9.885 | 10.009 | 10.041 | 9.812 | 10.360 |
| MRL-Up | NM_025912 | 2010011I20Rik | 9.471 | 8.967 | 8.233 | 9.619 | 9.699 | 9.529 | 9.345 | 9.334 | 9.300 | 8.899 | 9.039 | 8.870 | 9.183 | 8.845 | 8.152 | 8.259 | 7.989 | 8.352 | 8.415 |
| MRL-Up | NM_010589 | Jak3 | 9.723 | 7.806 | 7.181 | 9.646 | 9.923 | 9.820 | 9.680 | 9.493 | 9.777 | 7.772 | 7.953 | 7.645 | 7.990 | 7.668 | 7.228 | 7.128 | 6.984 | 7.223 | 7.340 |
| MRL-Up | NM_145450 | BC022687 | 11.774 | 10.722 | 9.898 | 11.742 | 11.702 | 11.811 | 11.868 | 11.771 | 11.749 | 10.728 | 10.747 | 10.627 | 10.511 | 10.995 | 9.944 | 9.842 | 9.772 | 9.535 | 10.396 |
| MRL-Up | NM_134248 | Havcr1 | 7.497 | 6.240 | 5.562 | 7.628 | 7.728 | 7.533 | 7.424 | 7.226 | 7.446 | 5.999 | 6.401 | 6.269 | 6.353 | 6.180 | 5.698 | 5.503 | 5.481 | 5.468 | 5.661 |
| MRL-Up | NM_053007 | Cntf | 11.271 | 10.463 | 9.697 | 11.027 | 11.238 | 11.522 | 11.706 | 11.178 | 10.953 | 10.355 | 10.500 | 10.488 | 10.498 | 10.472 | 9.743 | 9.953 | 9.781 | 9.356 | 9.649 |
| MRL-Up | NM_011035 | Pak1 | 11.175 | 10.410 | 8.930 | 11.122 | 11.465 | 11.278 | 11.065 | 10.996 | 11.126 | 10.550 | 10.509 | 10.273 | 10.456 | 10.262 | 9.202 | 8.687 | 8.686 | 9.261 | 8.812 |
| MRL-Up | XM_975103 | LOC665180 | 9.377 | 8.388 | 7.839 | 9.185 | 9.560 | 9.172 | 9.270 | 9.412 | 9.665 | 8.571 | 8.274 | 8.299 | 8.190 | 8.607 | 7.794 | 7.946 | 8.189 | 7.372 | 7.893 |
| MRL-Up | AK087705 | Mmp28 | 9.892 | 9.371 | 7.410 | 9.896 | 10.322 | 9.994 | 9.755 | 9.694 | 9.694 | 9.307 | 9.710 | 8.902 | 9.707 | 9.228 | 7.198 | 7.252 | 7.545 | 7.223 | 7.832 |
| MRL-Up | NM_130455 | Grin3b | 8.588 | 8.068 | 6.672 | 8.745 | 8.492 | 8.537 | 8.457 | 8.780 | 8.513 | 8.042 | 8.031 | 8.029 | 8.197 | 8.044 | 6.760 | 6.494 | 6.899 | 6.712 | 6.492 |
| MRL-Up | NM_009034 | Rbp2 | 12.561 | 11.403 | 10.263 | 12.473 | 12.766 | 12.496 | 12.426 | 12.311 | 12.893 | 11.401 | 11.510 | 11.400 | 11.437 | 11.268 | 10.019 | 10.895 | 10.088 | 9.938 | 10.375 |
| MRL-Up | NM_008499 | Lhx5 | 6.994 | 6.160 | 5.501 | 7.110 | 7.340 | 6.932 | 6.815 | 6.640 | 7.130 | 5.980 | 6.468 | 6.162 | 6.212 | 5.980 | 5.704 | 5.611 | 5.362 | 5.290 | 5.535 |
| MRL-Up | NM_026496 | Grhl2 | 8.292 | 7.742 | 6.958 | 8.248 | 8.784 | 8.225 | 8.090 | 7.966 | 8.440 | 7.596 | 8.043 | 7.657 | 7.915 | 7.498 | 7.006 | 7.348 | 6.733 | 6.658 | 7.047 |

Supplemental Table S1 (continued)

| Group | Genbank ID | GeneName | Mean MRL log2 intensity | Mean WT log2 intensity | Mean PAX6 log2 intensity | MRL_1 log2 intensity | MRL_2 log2 intensity | MRL_3 log2 intensity | MRL_4 log2 intensity | MRL_5 log2 intensity | MRL_6 log2 intensity | WT_1 log2 intensity | WT_2 log2 intensity | WT_3 log2 intensity | WT_4 log2 intensity | WT_5 log2 intensity | PAX6_1 log2 intensity | PAX6_2 log2 intensity | PAX6_3 log2 intensity | PAX6_4 log2 intensity | PAX6_5 log2 intensity |
|---|---|---|---|---|---|---|---|---|---|---|---|---|---|---|---|---|---|---|---|---|---|
| MRL-Up | NM_019808 | Pdlim5 | 12.319 | 11.053 | 10.497 | 12.251 | 12.462 | 12.437 | 12.324 | 12.149 | 12.291 | 11.082 | 11.218 | 11.066 | 11.034 | 10.866 | 10.839 | 10.746 | 10.429 | 9.951 | 10.519 |
| MRL-Up | NM_178848 | Sirt5 | 11.490 | 10.759 | 10.073 | 11.470 | 11.436 | 11.667 | 11.595 | 11.615 | 11.158 | 10.662 | 10.655 | 10.859 | 10.944 | 10.673 | 9.949 | 9.767 | 10.104 | 10.399 | 10.145 |
| MRL-Up | NM_133900 | Psph | 11.407 | 10.581 | 10.095 | 11.140 | 11.345 | 11.257 | 11.296 | 11.420 | 11.986 | 10.460 | 10.508 | 10.770 | 10.400 | 10.765 | 10.343 | 10.069 | 9.953 | 9.861 | 10.248 |
| MRL-Up | NM_007429 | Agtr2 | 11.711 | 10.753 | 8.881 | 11.794 | 12.052 | 11.848 | 11.883 | 11.136 | 11.550 | 10.560 | 10.872 | 10.711 | 10.914 | 10.710 | 8.701 | 9.229 | 8.727 | 8.561 | 9.185 |
| MRL-Up | NM_008290 | Hsd17b2 | 12.945 | 12.274 | 11.693 | 13.315 | 13.100 | 13.074 | 12.605 | 12.843 | 12.732 | 12.401 | 12.447 | 12.170 | 12.400 | 11.952 | 11.952 | 11.585 | 11.392 | 11.238 | 12.299 |
| MRL-Up | NM_008540 | Smad4 | 11.476 | 10.896 | 9.454 | 11.603 | 11.455 | 11.647 | 11.562 | 11.370 | 11.218 | 10.750 | 10.913 | 10.731 | 11.309 | 10.776 | 10.061 | 9.024 | 9.356 | 9.410 | 9.416 |
| MRL-Up | NM_153092 | Nupl2 | 8.521 | 7.974 | 7.419 | 8.538 | 8.710 | 8.607 | 8.611 | 8.272 | 8.386 | 7.881 | 8.033 | 7.980 | 8.089 | 7.887 | 7.285 | 7.308 | 7.258 | 7.612 | 7.633 |
| MRL-Up | NM_029809 | 2310014L17Rik | 7.567 | 6.672 | 6.096 | 7.653 | 7.777 | 7.644 | 7.540 | 7.486 | 7.299 | 6.445 | 6.739 | 6.525 | 6.961 | 6.692 | 6.191 | 6.156 | 6.112 | 5.807 | 6.215 |
| MRL-Up | NM_001003913 | Mars | 12.624 | 11.982 | 11.441 | 12.437 | 12.477 | 12.764 | 12.467 | 12.722 | 12.879 | 12.128 | 11.765 | 11.872 | 12.174 | 11.972 | 11.452 | 11.463 | 11.358 | 11.599 | 11.333 |
| MRL-Up | AV333862 | AI853839 | 6.822 | 6.069 | 5.432 | 6.880 | 6.757 | 6.839 | 7.070 | 6.660 | 6.723 | 5.834 | 6.366 | 6.138 | 6.153 | 5.852 | 5.386 | 5.344 | 5.797 | 5.369 | 5.265 |
| MRL-Up | NM_019955 | Ripk3 | 10.907 | 10.284 | 9.497 | 10.871 | 11.270 | 10.914 | 10.769 | 10.600 | 11.019 | 10.259 | 10.275 | 10.274 | 10.399 | 10.212 | 9.478 | 9.255 | 9.470 | 9.193 | 10.089 |
| MRL-Up | NM_134249 | Timd2 | 7.927 | 6.699 | 5.896 | 8.032 | 8.186 | 7.918 | 7.877 | 7.588 | 7.964 | 6.704 | 6.824 | 6.595 | 6.961 | 6.408 | 6.038 | 5.786 | 5.787 | 5.925 | 5.947 |
| MRL-Up | NM_172898 | Kirrel2 | 6.800 | 6.146 | 5.446 | 7.109 | 6.780 | 6.968 | 6.673 | 6.383 | 6.883 | 6.324 | 6.078 | 6.388 | 5.836 | 6.102 | 5.466 | 5.322 | 5.409 | 5.638 | 5.397 |
| MRL-Up | NM_198193 | Raet1e | 7.272 | 6.296 | 5.717 | 7.372 | 7.415 | 7.499 | 7.194 | 7.240 | 6.913 | 6.175 | 6.373 | 6.263 | 6.529 | 6.139 | 5.796 | 5.781 | 5.591 | 5.700 | 5.715 |
| MRL-Up | NM_199017 | 9230110C19Rik | 8.005 | 7.363 | 6.756 | 8.021 | 8.087 | 8.141 | 7.976 | 7.842 | 7.962 | 7.213 | 7.381 | 7.424 | 7.615 | 7.182 | 6.901 | 6.541 | 6.732 | 6.799 | 6.806 |
| MRL-Up | NM_134249 | Timd2 | 7.716 | 7.048 | 5.828 | 7.661 | 8.059 | 7.615 | 7.570 | 7.382 | 8.007 | 6.786 | 7.296 | 7.100 | 7.122 | 6.938 | 6.073 | 5.801 | 5.622 | 5.823 | 5.823 |
| MRL-Up | NM_028439 | 3110009E18Rik | 10.717 | 10.174 | 9.044 | 10.807 | 11.014 | 10.467 | 10.790 | 10.480 | 10.746 | 9.995 | 10.224 | 10.244 | 10.067 | 10.342 | 9.352 | 9.048 | 8.879 | 8.766 | 9.178 |
| MRL-Up | AK085462 | AK085462 | 8.830 | 8.289 | 7.228 | 8.908 | 9.029 | 8.904 | 8.741 | 8.643 | 8.752 | 8.333 | 8.342 | 8.347 | 8.449 | 7.972 | 7.258 | 7.101 | 7.058 | 7.147 | 7.575 |
| MRL-Up | AK084717 | Spon1 | 12.587 | 10.300 | 9.195 | 12.689 | 12.579 | 12.780 | 12.684 | 12.351 | 12.439 | 10.262 | 10.415 | 10.265 | 10.425 | 10.133 | 9.406 | 9.251 | 8.770 | 9.559 | 8.987 |
| MRL-Up | AV304517 | AV304517 | 8.203 | 7.638 | 7.008 | 8.142 | 8.104 | 7.945 | 8.263 | 8.149 | 8.612 | 7.502 | 7.612 | 7.685 | 7.572 | 7.819 | 7.028 | 6.905 | 6.998 | 7.174 | 6.936 |
| MRL-Up | BC100444 | 4930579G22Rik | 10.870 | 9.898 | 8.706 | 10.920 | 10.814 | 10.772 | 10.738 | 11.005 | 10.970 | 9.921 | 9.928 | 9.924 | 9.827 | 9.891 | 8.861 | 8.576 | 8.596 | 8.645 | 8.853 |
| MRL-Up | NM_198104 | Tcte3 | 7.868 | 7.372 | 5.777 | 7.869 | 8.076 | 8.049 | 7.734 | 7.655 | 7.826 | 7.238 | 7.437 | 7.349 | 7.576 | 7.262 | 5.883 | 5.564 | 5.882 | 5.638 | 5.920 |
| MRL-Up | NM_001003913 | Mars | 10.891 | 10.193 | 9.610 | 10.662 | 10.822 | 11.070 | 10.826 | 10.909 | 11.059 | 10.238 | 10.082 | 10.206 | 10.402 | 10.037 | 9.692 | 9.542 | 9.061 | 9.966 | 9.792 |
| MRL-Up | AK082372 | Ppp1r14c | 7.118 | 6.406 | 5.861 | 6.936 | 7.328 | 7.165 | 7.083 | 7.076 | 7.121 | 6.197 | 6.623 | 6.314 | 6.677 | 6.222 | 5.815 | 5.849 | 5.702 | 5.822 | 6.117 |
| MRL-Up | XM_984921 | EG237412 | 7.872 | 7.367 | 6.837 | 7.915 | 7.905 | 8.079 | 8.069 | 7.841 | 7.421 | 7.211 | 7.324 | 7.479 | 7.504 | 7.319 | 6.760 | 6.913 | 6.633 | 6.767 | 7.112 |
| MRL-Up | NM_001039552 | 9030025P20Rik | 9.198 | 8.531 | 7.608 | 9.151 | 9.177 | 9.286 | 9.409 | 9.125 | 9.043 | 8.410 | 8.638 | 8.491 | 8.752 | 8.363 | 7.663 | 7.386 | 7.399 | 7.867 | 7.723 |
| MRL-Up | NM_178610 | Krr1 | 9.774 | 8.899 | 8.309 | 9.829 | 9.842 | 9.849 | 9.895 | 9.703 | 9.527 | 8.694 | 8.895 | 9.102 | 9.033 | 8.770 | 8.825 | 8.143 | 8.172 | 8.211 | 8.193 |
| MRL-Up | NM_027495 | Tmem144 | 9.303 | 8.499 | 7.454 | 9.380 | 9.479 | 9.224 | 9.017 | 9.213 | 9.506 | 8.497 | 8.643 | 8.324 | 8.633 | 8.398 | 7.527 | 7.438 | 7.457 | 7.399 | 7.448 |
| MRL-Up | NM_007904 | Ednrb | 11.424 | 10.766 | 8.864 | 11.844 | 10.950 | 11.662 | 11.456 | 11.110 | 11.525 | 10.771 | 10.690 | 11.414 | 10.251 | 10.702 | 8.900 | 8.584 | 8.751 | 8.973 | 9.110 |
| MRL-Up | NM_027495 | Tmem144 | 8.653 | 8.132 | 7.037 | 8.773 | 8.833 | 8.667 | 8.576 | 8.497 | 8.573 | 7.943 | 8.201 | 8.161 | 8.340 | 8.013 | 7.261 | 7.075 | 6.790 | 6.932 | 7.128 |
| MRL-Up | XR_002830 | EG627488 | 10.249 | 9.705 | 8.934 | 10.198 | 10.470 | 10.183 | 10.111 | 10.045 | 10.491 | 9.441 | 9.785 | 9.702 | 9.796 | 9.802 | 9.183 | 8.915 | 8.929 | 8.764 | 8.877 |
| MRL-Up | NM_207675 | Cadm1 | 14.017 | 12.734 | 11.598 | 14.127 | 13.591 | 13.914 | 14.093 | 14.287 | 14.091 | 12.886 | 12.601 | 12.902 | 12.392 | 12.890 | 11.875 | 11.462 | 11.250 | 11.595 | 11.808 |
| MRL-Up | AK145337 | Chchd3 | 9.765 | 9.031 | 8.429 | 9.803 | 9.814 | 9.643 | 9.695 | 9.791 | 9.841 | 9.091 | 9.083 | 9.050 | 9.014 | 8.918 | 8.605 | 8.585 | 8.325 | 8.366 | 8.262 |
| MRL-Up | NM_177175 | A930001M12Rik | 7.154 | 6.569 | 5.437 | 7.168 | 7.331 | 7.045 | 7.150 | 7.191 | 7.040 | 6.416 | 6.489 | 6.661 | 6.400 | 6.878 | 5.432 | 5.434 | 5.367 | 5.366 | 5.588 |

Supplemental Table S1 (continued)

| Group | Genbank ID | GeneName | Mean MRL log2 intensity | Mean WT log2 intensity | Mean PAX6 log2 intensity | MRL_1 log2 intensity | MRL_2 log2 intensity | MRL_3 log2 intensity | MRL_4 log2 intensity | MRL_5 log2 intensity | MRL_6 log2 intensity | WT_1 log2 intensity | WT_2 log2 intensity | WT_3 log2 intensity | WT_4 log2 intensity | WT_5 log2 intensity | PAX6_1 log2 intensity | PAX6_2 log2 intensity | PAX6_3 log2 intensity | PAX6_4 log2 intensity | PAX6_5 log2 intensity |
|---|---|---|---|---|---|---|---|---|---|---|---|---|---|---|---|---|---|---|---|---|---|
| MRL-Up | NM_183024 | Raver2 | 11.257 | 10.652 | 7.388 | 11.412 | 11.418 | 11.424 | 11.201 | 11.131 | 10.954 | 10.375 | 10.924 | 10.780 | 10.872 | 10.310 | 7.588 | 7.102 | 7.289 | 7.693 | 7.265 |
| MRL-Up | NM_019808 | Pdlim5 | 11.973 | 10.842 | 10.154 | 11.906 | 12.188 | 12.098 | 12.012 | 11.720 | 11.912 | 10.899 | 11.080 | 10.822 | 10.793 | 10.618 | 10.470 | 10.429 | 10.078 | 9.628 | 10.167 |
| MRL-Up | AK043775 | B230206I08Rik | 7.105 | 6.371 | 5.814 | 7.171 | 7.345 | 7.158 | 6.968 | 7.098 | 6.890 | 6.432 | 6.491 | 6.216 | 6.286 | 6.433 | 5.940 | 5.694 | 5.828 | 5.841 | 5.765 |
| MRL-Up | BC032927 | Hist1h1e | 8.338 | 7.427 | 6.313 | 8.226 | 8.620 | 8.209 | 8.271 | 8.373 | 8.326 | 7.374 | 7.350 | 7.501 | 7.213 | 7.697 | 6.397 | 6.177 | 6.223 | 6.224 | 6.544 |
| MRL-Up | AK080551 | AK080551 | 8.872 | 8.386 | 7.495 | 8.552 | 8.902 | 8.488 | 8.751 | 9.322 | 9.218 | 8.502 | 8.125 | 8.372 | 8.093 | 8.838 | 7.635 | 7.511 | 7.549 | 7.204 | 7.575 |
| MRL-Up | NM_001025606 | Tmem171 | 7.913 | 7.369 | 6.229 | 7.752 | 8.177 | 7.945 | 7.953 | 7.454 | 8.196 | 7.364 | 7.707 | 7.092 | 7.370 | 7.312 | 6.203 | 6.277 | 5.878 | 6.291 | 6.495 |
| MRL-Up | BC063330 | 1200003I07Rik | 7.896 | 7.293 | 6.807 | 7.904 | 8.267 | 7.780 | 7.857 | 7.738 | 7.828 | 7.223 | 7.462 | 7.363 | 7.349 | 7.068 | 7.125 | 6.836 | 6.696 | 6.797 | 6.584 |

Supplementary Table S2

| Group | Genbank ID | GeneName | Mean MRL log2 intensity | Mean WT log2 intensity | Mean PAX6 log2 intensity | MRL_1 log2 intensity | MRL_2 log2 intensity | MRL_3 log2 intensity | MRL_4 log2 intensity | MRL_5 log2 intensity | MRL_6 log2 intensity | WT_1 log2 intensity | WT_2 log2 intensity | WT_3 log2 intensity | WT_4 log2 intensity | WT_5 log2 intensity | PAX6_1 log2 intensity | PAX6_2 log2 intensity | PAX6_3 log2 intensity | PAX6_4 log2 intensity | PAX6_5 log2 intensity |
|---|---|---|---|---|---|---|---|---|---|---|---|---|---|---|---|---|---|---|---|---|---|
| Pax6-Up | NM_033080 | Nudt19 | 11.650 | 12.268 | 13.813 | 11.587 | 11.697 | 11.847 | 11.724 | 11.551 | 11.492 | 12.357 | 12.233 | 12.289 | 12.408 | 12.053 | 13.757 | 13.930 | 14.003 | 13.797 | 13.578 |
| Pax6-Up | NM_019707 | Cdh13 | 8.788 | 9.636 | 11.359 | 8.634 | 8.950 | 8.824 | 9.056 | 8.797 | 8.467 | 9.773 | 9.587 | 9.678 | 9.778 | 9.367 | 10.941 | 11.307 | 11.751 | 11.206 | 11.587 |
| Pax6-Up | NM_009402 | Pglyrp1 | 13.253 | 13.781 | 15.456 | 12.787 | 13.558 | 13.381 | 13.663 | 12.834 | 13.295 | 13.746 | 13.750 | 13.935 | 13.712 | 13.762 | 15.231 | 15.647 | 15.790 | 15.035 | 15.577 |
| Pax6-Up | NM_001080941 | 2810487A22Rik | 7.377 | 9.353 | 9.952 | 7.435 | 7.515 | 7.302 | 7.458 | 7.217 | 7.336 | 9.299 | 9.501 | 9.596 | 9.311 | 9.056 | 10.044 | 10.250 | 9.838 | 9.681 | 9.949 |
| Pax6-Up | NM_008529 | Ly6e | 11.228 | 11.874 | 14.049 | 11.020 | 11.411 | 11.426 | 11.606 | 10.505 | 11.401 | 12.091 | 11.935 | 11.738 | 11.920 | 11.686 | 13.624 | 14.411 | 14.143 | 14.021 | 14.048 |
| Pax6-Up | NM_199197 | 1110032A13Rik | 9.421 | 10.072 | 10.728 | 9.446 | 9.595 | 9.491 | 9.330 | 9.281 | 9.385 | 10.180 | 10.219 | 9.890 | 10.166 | 9.903 | 10.387 | 10.788 | 10.921 | 10.748 | 10.794 |
| Pax6-Up | NM_010217 | Ctgf | 11.771 | 12.267 | 15.763 | 11.670 | 11.915 | 11.671 | 12.485 | 11.750 | 11.136 | 12.015 | 12.025 | 12.987 | 11.901 | 12.408 | 15.908 | 15.686 | 15.905 | 15.467 | 15.850 |
| Pax6-Up | NM_008035 | Folr2 | 8.707 | 9.325 | 11.887 | 8.362 | 8.765 | 8.827 | 9.160 | 8.416 | 8.712 | 9.422 | 9.487 | 9.444 | 9.371 | 8.900 | 11.874 | 12.576 | 11.725 | 11.766 | 11.492 |
| Pax6-Up | NM_011926 | Ceacam1 | 9.895 | 10.771 | 12.736 | 9.851 | 10.355 | 9.896 | 10.059 | 9.339 | 9.870 | 10.727 | 10.723 | 10.618 | 11.020 | 10.764 | 12.620 | 13.364 | 12.299 | 12.499 | 12.898 |
| Pax6-Up | NM_027219 | Cdc42ep1 | 7.931 | 8.540 | 9.182 | 8.003 | 8.141 | 8.125 | 7.863 | 7.830 | 7.625 | 8.708 | 8.685 | 8.505 | 8.616 | 8.186 | 9.208 | 9.150 | 9.489 | 9.083 | 8.981 |
| Pax6-Up | NM_028968 | Ifitm7 | 13.078 | 13.600 | 14.575 | 13.012 | 13.183 | 13.390 | 13.173 | 12.519 | 13.194 | 13.777 | 13.579 | 13.364 | 13.750 | 13.528 | 14.225 | 14.644 | 14.537 | 14.648 | 14.820 |
| Pax6-Up | M69070 | H2-K1 | 13.431 | 14.134 | 14.621 | 13.210 | 13.414 | 13.840 | 13.470 | 13.306 | 13.347 | 14.131 | 14.204 | 14.112 | 14.214 | 14.008 | 14.400 | 15.167 | 14.172 | 15.240 | 14.127 |
| Pax6-Up | D86232 | Ly6c | 12.409 | 13.422 | 15.019 | 12.196 | 12.349 | 12.694 | 13.016 | 11.848 | 12.352 | 13.672 | 13.499 | 13.109 | 13.529 | 13.299 | 14.398 | 15.465 | 14.894 | 15.203 | 15.136 |
| Pax6-Up | NM_001001892 | H2-K1 | 14.185 | 14.881 | 15.398 | 14.039 | 14.153 | 14.558 | 14.186 | 14.099 | 14.073 | 14.873 | 14.931 | 14.915 | 14.955 | 14.731 | 15.115 | 15.846 | 15.049 | 16.081 | 14.898 |
| Pax6-Up | M18187 | Lst1 | 9.190 | 9.896 | 10.548 | 8.925 | 9.297 | 9.191 | 9.366 | 8.907 | 9.452 | 9.855 | 10.039 | 10.039 | 9.877 | 9.667 | 10.339 | 10.525 | 11.063 | 10.422 | 10.388 |
| Pax6-Up | NM_008609 | Mmp15 | 8.454 | 9.296 | 11.157 | 8.374 | 8.897 | 8.427 | 8.540 | 8.291 | 8.193 | 9.434 | 9.312 | 9.241 | 9.377 | 9.117 | 10.856 | 11.509 | 11.269 | 10.944 | 11.208 |
| Pax6-Up | NM_010581 | Cd47 | 11.722 | 12.251 | 12.841 | 11.824 | 11.923 | 11.795 | 11.877 | 11.665 | 11.247 | 12.142 | 12.417 | 12.188 | 12.340 | 12.169 | 12.849 | 12.902 | 12.466 | 12.882 | 13.104 |
| Pax6-Up | NM_008080 | B4galnt1 | 7.881 | 9.438 | 10.389 | 7.673 | 7.850 | 7.991 | 8.121 | 7.673 | 7.977 | 9.631 | 9.503 | 9.202 | 9.733 | 9.121 | 10.591 | 10.504 | 10.659 | 9.980 | 10.212 |
| Pax6-Up | XM_283804 | 9830001H06Rik | 5.995 | 6.648 | 7.441 | 5.969 | 6.064 | 5.957 | 6.006 | 5.885 | 6.092 | 6.697 | 6.720 | 6.678 | 6.668 | 6.477 | 7.391 | 7.528 | 7.531 | 7.521 | 7.235 |
| Pax6-Up | AF024519 | Tsc22d3 | 12.294 | 12.982 | 15.494 | 12.536 | 12.377 | 12.199 | 12.163 | 12.111 | 12.380 | 12.816 | 13.013 | 13.096 | 12.887 | 13.096 | 15.451 | 15.571 | 15.965 | 15.651 | 14.833 |
| Pax6-Up | NM_010286 | Tsc22d3 | 12.004 | 12.697 | 15.015 | 12.301 | 12.077 | 11.925 | 11.878 | 11.810 | 12.034 | 12.508 | 12.681 | 12.859 | 12.668 | 12.769 | 15.067 | 15.175 | 15.437 | 15.067 | 14.326 |
| Pax6-Up | NM_009950 | Cradd | 6.079 | 10.341 | 10.908 | 6.050 | 6.047 | 6.130 | 6.096 | 6.044 | 6.111 | 10.323 | 10.331 | 10.460 | 10.204 | 10.388 | 10.584 | 11.152 | 11.157 | 10.738 | 10.906 |
| Pax6-Up | AK030069 | Olfml2a | 6.358 | 7.243 | 8.738 | 6.208 | 6.400 | 6.692 | 6.511 | 6.373 | 5.964 | 7.259 | 7.266 | 7.200 | 7.625 | 6.867 | 8.720 | 8.270 | 9.137 | 8.668 | 8.897 |
| Pax6-Up | NM_207666 | Egfl9 | 9.315 | 10.420 | 12.008 | 9.558 | 9.379 | 9.276 | 9.215 | 9.209 | 9.253 | 10.414 | 10.403 | 10.574 | 10.569 | 10.141 | 11.819 | 11.956 | 11.965 | 11.943 | 12.358 |
| Pax6-Up | NM_011878 | Tiam2 | 9.764 | 10.307 | 10.971 | 9.670 | 9.853 | 9.937 | 9.892 | 9.405 | 9.828 | 10.107 | 10.390 | 10.490 | 10.486 | 10.063 | 11.250 | 11.089 | 10.818 | 10.983 | 10.717 |
| Pax6-Up | NM_172935 | Amdhd2 | 11.211 | 12.009 | 12.749 | 11.178 | 11.203 | 11.210 | 11.219 | 11.399 | 11.055 | 12.148 | 11.998 | 12.029 | 12.059 | 11.810 | 12.391 | 12.890 | 13.155 | 12.606 | 12.701 |
| Pax6-Up | NM_013705 | Zfp30 | 7.641 | 8.162 | 8.901 | 7.605 | 7.359 | 7.601 | 7.987 | 7.649 | 7.647 | 7.925 | 8.144 | 8.467 | 7.940 | 8.336 | 8.705 | 8.955 | 9.120 | 8.893 | 8.832 |
| Pax6-Up | NM_029646 | 2010004A03Rik | 9.763 | 10.264 | 12.711 | 9.864 | 9.518 | 9.937 | 9.716 | 9.733 | 9.811 | 10.440 | 9.997 | 10.401 | 10.277 | 10.206 | 12.325 | 12.809 | 13.292 | 12.606 | 12.521 |
| Pax6-Up | NM_010387 | H2-DMb1 | 9.468 | 10.007 | 11.228 | 9.498 | 9.618 | 9.491 | 9.582 | 9.187 | 9.433 | 10.070 | 10.144 | 10.152 | 9.993 | 9.674 | 11.057 | 11.013 | 11.268 | 11.584 | 11.219 |
| Pax6-Up | NM_009825 | Serpinh1 | 13.994 | 14.487 | 15.516 | 14.052 | 13.808 | 13.893 | 14.011 | 14.025 | 14.175 | 14.606 | 14.441 | 14.601 | 14.494 | 14.293 | 16.020 | 15.240 | 15.831 | 15.605 | 14.883 |
| Pax6-Up | NM_019631 | Tmem45a | 10.163 | 12.256 | 12.981 | 10.174 | 10.533 | 10.107 | 10.280 | 9.777 | 10.110 | 12.286 | 12.256 | 12.217 | 12.355 | 12.167 | 13.127 | 12.978 | 12.989 | 12.534 | 13.278 |
| Pax6-Up | NM_008802 | Pde7a | 10.089 | 10.702 | 11.612 | 10.041 | 10.229 | 10.110 | 10.176 | 9.881 | 10.099 | 10.673 | 10.743 | 10.800 | 10.724 | 10.572 | 11.749 | 11.741 | 11.460 | 11.333 | 11.775 |
| Pax6-Up | NM_011385 | Ski | 10.720 | 11.373 | 12.225 | 10.931 | 10.816 | 10.795 | 10.486 | 10.688 | 10.606 | 11.648 | 11.421 | 11.158 | 11.516 | 11.123 | 12.018 | 12.181 | 12.777 | 12.420 | 11.728 |
| Pax6-Up | AK019508 | Gpr115 | 9.913 | 10.802 | 11.549 | 10.106 | 10.250 | 9.917 | 9.848 | 9.557 | 9.801 | 10.999 | 11.005 | 10.810 | 10.757 | 10.441 | 11.497 | 11.785 | 11.105 | 11.500 | 11.857 |
| Pax6-Up | NM_011723 | Xdh | 8.657 | 10.961 | 12.263 | 8.592 | 8.733 | 9.176 | 8.402 | 8.251 | 8.789 | 10.898 | 10.944 | 11.010 | 11.109 | 10.844 | 11.856 | 12.641 | 12.263 | 12.366 | 12.189 |
| Pax6-Up | NM_009726 | Atp7a | 9.636 | 10.132 | 10.670 | 9.834 | 9.733 | 9.785 | 9.450 | 9.566 | 9.447 | 10.223 | 10.074 | 10.156 | 10.068 | 10.138 | 10.603 | 10.098 | 11.035 | 11.017 | 10.597 |

Supplementary Table S2 (continued)

| Group | Genbank ID | GeneName | Mean MRL log2 intensity | Mean WT log2 intensity | Mean PAX6 log2 intensity | MRL_1 log2 intensity | MRL_2 log2 intensity | MRL_3 log2 intensity | MRL_4 log2 intensity | MRL_5 log2 intensity | MRL_6 log2 intensity | WT_1 log2 intensity | WT_2 log2 intensity | WT_3 log2 intensity | WT_4 log2 intensity | WT_5 log2 intensity | PAX6_1 log2 intensity | PAX6_2 log2 intensity | PAX6_3 log2 intensity | PAX6_4 log2 intensity | PAX6_5 log2 intensity |
|---|---|---|---|---|---|---|---|---|---|---|---|---|---|---|---|---|---|---|---|---|---|
| Pax6-Up | TC1620752 | TC1620752 | 9.532 | 10.050 | 10.630 | 9.541 | 9.883 | 9.709 | 9.526 | 9.204 | 9.330 | 10.126 | 10.221 | 9.948 | 10.262 | 9.695 | 10.168 | 10.267 | 10.871 | 11.023 | 10.824 |
| Pax6-Up | BC005527 | Enpp3 | 11.144 | 12.593 | 13.105 | 11.130 | 11.304 | 11.175 | 11.423 | 10.697 | 11.133 | 12.515 | 12.660 | 12.642 | 12.962 | 12.184 | 13.159 | 13.394 | 12.577 | 13.027 | 13.368 |
| Pax6-Up | NM_007985 | Fancc | 8.057 | 8.943 | 9.926 | 7.958 | 8.245 | 7.885 | 8.076 | 7.931 | 8.248 | 8.842 | 9.038 | 9.003 | 9.047 | 8.783 | 9.855 | 10.199 | 10.185 | 9.785 | 9.605 |
| Pax6-Up | NM_012057 | Irf5 | 9.054 | 9.664 | 10.379 | 9.225 | 9.408 | 9.230 | 9.027 | 8.667 | 8.768 | 9.646 | 9.777 | 9.667 | 9.812 | 9.417 | 10.155 | 10.598 | 10.223 | 10.324 | 10.593 |
| Pax6-Up | NM_145078 | 2610305D13Rik | 5.310 | 7.021 | 8.911 | 5.446 | 5.495 | 5.289 | 5.241 | 5.275 | 5.114 | 6.724 | 7.089 | 7.110 | 7.235 | 6.947 | 8.544 | 9.300 | 9.119 | 9.037 | 8.555 |
| Pax6-Up | NM_010592 | Jund1 | 14.280 | 14.785 | 15.657 | 14.182 | 14.210 | 14.308 | 14.335 | 14.301 | 14.345 | 14.755 | 14.873 | 14.966 | 14.754 | 14.578 | 16.466 | 15.599 | 15.927 | 15.282 | 15.013 |
| Pax6-Up | NM_023158 | Cxcl16 | 9.177 | 9.839 | 10.717 | 9.207 | 9.290 | 9.257 | 9.230 | 8.799 | 9.277 | 10.059 | 9.970 | 9.877 | 9.744 | 9.548 | 10.781 | 10.975 | 10.179 | 10.982 | 10.665 |
| Pax6-Up | NM_054094 | Acsm1 | 10.980 | 11.754 | 12.996 | 11.174 | 11.211 | 11.080 | 10.971 | 10.460 | 10.985 | 12.159 | 12.179 | 11.708 | 11.678 | 11.046 | 12.824 | 13.443 | 12.403 | 13.002 | 13.308 |
| Pax6-Up | AK017076 | 4933436C20Rik | 8.675 | 9.657 | 10.271 | 8.967 | 8.812 | 8.775 | 8.375 | 8.436 | 8.687 | 9.763 | 9.830 | 9.562 | 9.790 | 9.339 | 9.803 | 10.412 | 10.180 | 10.574 | 10.383 |
| Pax6-Up | NM_010330 | Emb | 11.329 | 11.883 | 13.944 | 11.246 | 11.435 | 11.231 | 11.688 | 11.143 | 11.231 | 12.010 | 12.155 | 11.949 | 12.077 | 11.225 | 13.853 | 13.733 | 14.344 | 13.968 | 13.822 |
| Pax6-Up | NM_020498 | Ly6i | 7.764 | 8.346 | 9.669 | 7.612 | 7.709 | 7.897 | 7.960 | 7.645 | 7.760 | 8.573 | 8.489 | 8.021 | 8.613 | 8.036 | 9.578 | 9.525 | 9.292 | 9.965 | 9.984 |
| Pax6-Up | NM_011408 | Slfn2 | 7.320 | 8.794 | 9.772 | 7.235 | 7.504 | 7.621 | 7.300 | 7.041 | 7.219 | 8.615 | 8.869 | 8.812 | 8.746 | 8.930 | 9.652 | 9.917 | 9.799 | 10.081 | 9.411 |
| Pax6-Up | NM_022995 | Tmepai | 8.640 | 9.293 | 10.329 | 8.610 | 8.726 | 8.737 | 8.299 | 8.879 | 8.588 | 9.374 | 9.113 | 9.389 | 9.178 | 9.408 | 10.001 | 9.970 | 10.388 | 10.686 | 10.600 |
| Pax6-Up | NM_011803 | Klf6 | 12.604 | 13.175 | 13.883 | 12.518 | 12.924 | 12.662 | 12.610 | 12.467 | 12.444 | 12.902 | 13.349 | 13.301 | 13.261 | 13.062 | 14.396 | 14.030 | 13.683 | 13.467 | 13.839 |
| Pax6-Up | NM_013605 | Muc1 | 6.580 | 7.237 | 9.690 | 6.452 | 6.728 | 6.643 | 6.890 | 6.217 | 6.547 | 7.480 | 7.231 | 7.431 | 7.277 | 6.766 | 9.509 | 9.919 | 9.232 | 10.164 | 9.627 |
| Pax6-Up | NM_145978 | Pdlim2 | 12.447 | 13.022 | 13.755 | 12.441 | 12.627 | 12.487 | 12.406 | 12.233 | 12.487 | 13.088 | 12.946 | 13.034 | 13.124 | 12.917 | 13.536 | 13.695 | 13.938 | 13.621 | 13.984 |
| Pax6-Up | NM_008469 | Krt15 | 15.726 | 17.047 | 17.621 | 15.389 | 16.173 | 15.197 | 16.631 | 15.261 | 15.706 | 17.326 | 17.045 | 16.992 | 17.082 | 16.792 | 17.356 | 17.761 | 17.636 | 17.268 | 18.084 |
| Pax6-Up | NM_138684 | Wfdc12 | 6.154 | 6.712 | 7.375 | 6.179 | 6.024 | 6.201 | 6.222 | 6.219 | 6.078 | 6.983 | 6.465 | 6.691 | 6.681 | 6.738 | 7.753 | 6.932 | 7.415 | 7.336 | 7.440 |
| Pax6-Up | NM_153081 | Slc16a11 | 13.427 | 14.613 | 16.321 | 13.134 | 13.372 | 13.462 | 13.815 | 13.408 | 13.368 | 14.779 | 14.669 | 14.628 | 14.784 | 14.207 | 16.048 | 16.392 | 16.471 | 16.435 | 16.261 |
| Pax6-Up | AK051559 | AK051559 | 6.239 | 6.862 | 7.501 | 6.184 | 6.234 | 6.347 | 6.254 | 6.222 | 6.194 | 6.869 | 6.729 | 6.952 | 6.927 | 6.835 | 7.210 | 7.446 | 7.476 | 7.594 | 7.779 |
| Pax6-Up | NM_001077495 | Pik3r1 | 9.699 | 10.260 | 10.790 | 9.779 | 9.752 | 9.818 | 9.640 | 9.628 | 9.575 | 10.226 | 10.355 | 10.253 | 10.328 | 10.136 | 10.942 | 10.562 | 11.185 | 10.865 | 10.397 |
| Pax6-Up | AK035520 | EG329217 | 5.286 | 6.061 | 7.542 | 5.278 | 5.331 | 5.294 | 5.292 | 5.265 | 5.253 | 5.949 | 6.062 | 6.055 | 6.339 | 5.897 | 7.416 | 7.975 | 7.471 | 7.536 | 7.312 |
| Pax6-Up | AK031663 | Nudt6 | 7.730 | 8.220 | 10.138 | 7.714 | 7.759 | 7.721 | 8.140 | 7.439 | 7.605 | 8.185 | 8.074 | 8.302 | 8.383 | 8.154 | 10.127 | 10.088 | 10.498 | 10.058 | 9.919 |
| Pax6-Up | AK162987 | AK162987 | 6.689 | 7.598 | 8.920 | 6.503 | 6.892 | 6.656 | 6.966 | 6.573 | 6.546 | 7.473 | 7.718 | 7.730 | 7.697 | 7.373 | 9.247 | 8.712 | 9.022 | 8.985 | 8.632 |
| Pax6-Up | NM_007717 | Cmah | 10.004 | 10.827 | 12.205 | 10.087 | 10.248 | 10.069 | 10.410 | 9.393 | 9.817 | 11.113 | 10.802 | 10.757 | 11.049 | 10.413 | 12.422 | 12.479 | 11.776 | 12.485 | 11.864 |
| Pax6-Up | NM_011926 | Ceacam1 | 7.515 | 8.091 | 9.593 | 7.515 | 7.814 | 7.658 | 7.504 | 7.075 | 7.524 | 8.097 | 8.329 | 7.896 | 8.326 | 7.809 | 9.718 | 9.737 | 8.990 | 9.539 | 9.983 |
| Pax6-Up | BC004616 | Gusb | 6.921 | 8.718 | 9.384 | 6.956 | 7.167 | 6.942 | 6.816 | 6.836 | 6.807 | 8.763 | 8.762 | 8.676 | 8.819 | 8.570 | 9.149 | 9.221 | 9.797 | 9.469 | 9.283 |
| Pax6-Up | AK004111 | 1110035H17Rik | 6.373 | 6.923 | 7.472 | 6.437 | 6.287 | 6.223 | 6.289 | 6.484 | 6.519 | 6.948 | 6.960 | 6.963 | 6.832 | 6.912 | 7.590 | 7.461 | 7.784 | 7.185 | 7.341 |
| Pax6-Up | NM_009285 | Stc1 | 6.850 | 7.347 | 10.150 | 6.753 | 7.095 | 6.820 | 7.230 | 6.599 | 6.600 | 7.622 | 7.051 | 7.159 | 7.586 | 7.315 | 10.128 | 10.233 | 10.469 | 10.232 | 9.686 |
| Pax6-Up | AK028720 | 9530091C08Rik | 9.971 | 11.983 | 12.835 | 10.255 | 10.082 | 9.900 | 9.509 | 10.023 | 10.056 | 12.209 | 11.822 | 11.776 | 11.948 | 12.159 | 12.575 | 12.775 | 13.396 | 12.965 | 12.464 |
| Pax6-Up | NM_009982 | Ctsc | 10.520 | 11.016 | 12.072 | 10.575 | 10.460 | 10.622 | 10.621 | 10.369 | 10.473 | 10.956 | 10.996 | 10.981 | 11.187 | 10.961 | 12.510 | 11.961 | 12.127 | 12.078 | 11.685 |
| Pax6-Up | XM_283804 | 9830001H06Rik | 6.833 | 8.201 | 9.722 | 6.695 | 6.740 | 6.831 | 6.910 | 6.850 | 6.973 | 8.240 | 8.111 | 8.391 | 8.441 | 7.822 | 9.618 | 9.363 | 10.002 | 10.160 | 9.469 |
| Pax6-Up | NM_018861 | Slc1a4 | 9.271 | 9.765 | 10.601 | 8.958 | 9.618 | 9.422 | 9.304 | 8.962 | 9.360 | 9.700 | 9.528 | 9.940 | 9.901 | 9.756 | 10.685 | 10.672 | 10.675 | 10.431 | 10.542 |
| Pax6-Up | NM_027172 | 2310046K01Rik | 8.358 | 9.122 | 9.679 | 8.263 | 8.594 | 8.463 | 8.341 | 8.203 | 8.285 | 9.240 | 9.181 | 8.856 | 9.396 | 8.938 | 9.586 | 10.068 | 9.569 | 9.700 | 9.471 |
| Pax6-Up | NM_010851 | Myd88 | 10.782 | 11.831 | 12.404 | 10.764 | 11.074 | 10.853 | 10.766 | 10.478 | 10.759 | 11.894 | 11.921 | 11.654 | 11.875 | 11.810 | 12.384 | 12.516 | 12.398 | 12.334 | 12.388 |
| Pax6-Up | NM_172775 | Plxnb1 | 11.097 | 11.616 | 12.113 | 11.263 | 11.196 | 11.192 | 11.003 | 11.077 | 10.849 | 11.708 | 11.778 | 11.520 | 11.631 | 11.445 | 11.726 | 12.207 | 11.729 | 12.246 | 12.657 |

Supplementary Table S2 (continued)

| Group | Genbank ID | GeneName | Mean MRL log2 intensity | Mean WT log2 intensity | Mean PAX6 log2 intensity | MRL_1 log2 intensity | MRL_2 log2 intensity | MRL_3 log2 intensity | MRL_4 log2 intensity | MRL_5 log2 intensity | MRL_6 log2 intensity | WT_1 log2 intensity | WT_2 log2 intensity | WT_3 log2 intensity | WT_4 log2 intensity | WT_5 log2 intensity | PAX6_1 log2 intensity | PAX6_2 log2 intensity | PAX6_3 log2 intensity | PAX6_4 log2 intensity | PAX6_5 log2 intensity |
|---|---|---|---|---|---|---|---|---|---|---|---|---|---|---|---|---|---|---|---|---|---|
| Pax6-Up | AK048349 | AK048349 | 10.374 | 11.017 | 11.704 | 10.601 | 10.519 | 10.150 | 10.280 | 10.255 | 10.442 | 10.947 | 11.056 | 10.994 | 11.135 | 10.955 | 12.157 | 11.737 | 11.945 | 11.372 | 11.309 |
| Pax6-Up | NM_001039959 | Ahnak | 10.546 | 11.302 | 11.865 | 10.363 | 10.785 | 10.612 | 10.461 | 10.304 | 10.751 | 11.297 | 11.331 | 11.254 | 11.236 | 11.392 | 11.896 | 11.936 | 11.892 | 11.613 | 11.986 |
| Pax6-Up | NM_145973 | Ell3 | 7.886 | 8.663 | 10.286 | 8.003 | 8.139 | 7.767 | 7.854 | 7.598 | 7.958 | 8.579 | 8.814 | 8.640 | 8.764 | 8.517 | 10.198 | 10.194 | 10.213 | 10.196 | 10.630 |
| Pax6-Up | NM_011027 | P2rx7 | 9.439 | 10.230 | 11.485 | 9.320 | 9.482 | 9.603 | 9.462 | 9.329 | 9.438 | 10.178 | 10.439 | 10.372 | 10.366 | 9.792 | 11.370 | 11.630 | 11.484 | 11.573 | 11.366 |
| Pax6-Up | NM_010574 | Irx2 | 8.956 | 9.464 | 9.984 | 9.143 | 9.393 | 8.870 | 8.860 | 8.484 | 8.984 | 9.222 | 9.636 | 9.620 | 9.632 | 9.212 | 10.052 | 9.737 | 10.110 | 10.319 | 9.702 |
| Pax6-Up | ENSMUST00000091560 | ENSMUST00000091560 | 11.400 | 11.921 | 12.857 | 11.410 | 11.596 | 11.442 | 11.523 | 11.124 | 11.303 | 11.932 | 12.045 | 11.935 | 12.004 | 11.687 | 12.445 | 12.790 | 12.883 | 12.976 | 13.191 |
| Pax6-Up | BC039980 | Megf6 | 8.515 | 9.122 | 11.335 | 8.588 | 8.444 | 8.718 | 8.713 | 8.178 | 8.452 | 9.202 | 9.281 | 9.207 | 9.375 | 8.548 | 11.087 | 10.798 | 11.397 | 11.511 | 11.881 |
| Pax6-Up | NM_018851 | Samhd1 | 8.557 | 10.046 | 11.549 | 8.464 | 8.579 | 8.579 | 8.788 | 8.484 | 8.450 | 9.955 | 10.125 | 9.899 | 10.170 | 10.081 | 11.539 | 11.488 | 11.042 | 11.632 | 12.046 |
| Pax6-Up | NM_021542 | Kcnk5 | 9.631 | 10.126 | 10.622 | 9.842 | 9.777 | 9.998 | 9.429 | 9.231 | 9.507 | 10.359 | 10.224 | 9.965 | 10.318 | 9.762 | 10.630 | 10.789 | 10.277 | 10.488 | 10.924 |
| Pax6-Up | NM_009829 | Ccnd2 | 12.177 | 12.751 | 13.614 | 12.307 | 12.241 | 12.193 | 12.206 | 12.049 | 12.068 | 12.996 | 12.852 | 12.522 | 12.728 | 12.654 | 13.266 | 13.603 | 14.086 | 13.624 | 13.491 |
| Pax6-Up | NM_026820 | Ifitm1 | 13.695 | 14.469 | 15.178 | 13.556 | 13.886 | 13.786 | 13.936 | 13.134 | 13.873 | 14.637 | 14.485 | 14.296 | 14.510 | 14.419 | 14.690 | 15.451 | 15.104 | 15.059 | 15.588 |
| Pax6-Up | NM_027216 | Slc39a11 | 10.311 | 10.954 | 11.499 | 10.292 | 10.500 | 10.231 | 10.315 | 9.986 | 10.541 | 10.841 | 10.924 | 10.988 | 11.098 | 10.918 | 11.643 | 11.634 | 11.321 | 11.426 | 11.469 |
| Pax6-Up | NM_031159 | Apobec1 | 6.030 | 6.578 | 7.417 | 5.866 | 6.159 | 6.116 | 6.046 | 5.939 | 6.054 | 6.514 | 6.733 | 6.469 | 6.715 | 6.459 | 7.433 | 7.517 | 7.467 | 7.319 | 7.349 |
| Pax6-Up | AK040647 | AK040647 | 6.571 | 7.095 | 7.582 | 6.629 | 6.479 | 6.596 | 6.555 | 6.693 | 6.474 | 7.113 | 7.056 | 7.153 | 7.342 | 6.814 | 7.606 | 7.707 | 7.555 | 7.694 | 7.349 |
| Pax6-Up | BC080858 | Kbtbd11 | 11.257 | 11.783 | 13.659 | 11.186 | 10.873 | 11.305 | 11.539 | 11.378 | 11.259 | 12.150 | 11.731 | 11.694 | 11.625 | 11.716 | 13.353 | 14.114 | 13.648 | 13.994 | 13.187 |
| Pax6-Up | NM_027219 | Cdc42ep1 | 10.430 | 11.640 | 12.222 | 10.364 | 10.393 | 10.411 | 10.492 | 10.369 | 10.550 | 11.618 | 11.744 | 11.730 | 11.599 | 11.506 | 12.442 | 12.388 | 12.357 | 12.030 | 11.894 |
| Pax6-Up | NM_028079 | 2010111I01Rik | 10.653 | 11.384 | 12.114 | 10.762 | 10.907 | 10.573 | 10.785 | 10.142 | 10.748 | 11.063 | 11.425 | 11.787 | 11.390 | 11.253 | 11.974 | 11.992 | 12.064 | 11.971 | 12.568 |
| Pax6-Up | NM_010286 | Tsc22d3 | 10.536 | 11.202 | 13.030 | 10.865 | 10.558 | 10.572 | 10.485 | 10.358 | 10.381 | 10.987 | 11.169 | 11.329 | 11.371 | 11.154 | 13.403 | 12.920 | 12.630 | 13.519 | 12.680 |
| Pax6-Up | NM_011809 | Ets2 | 11.962 | 12.527 | 13.451 | 12.215 | 12.203 | 12.020 | 11.958 | 11.612 | 11.766 | 12.545 | 12.563 | 12.476 | 12.656 | 12.396 | 13.552 | 13.512 | 13.326 | 13.391 | 13.473 |
| Pax6-Up | AK080874 | Depdc6 | 7.404 | 8.719 | 11.052 | 7.312 | 7.434 | 7.429 | 7.396 | 7.421 | 7.433 | 8.900 | 8.379 | 9.005 | 8.366 | 8.943 | 10.820 | 11.004 | 11.600 | 11.171 | 10.664 |
| Pax6-Up | CN524910 | CN524910 | 8.526 | 9.285 | 10.248 | 8.449 | 8.694 | 8.564 | 8.660 | 8.417 | 8.370 | 9.272 | 9.262 | 9.370 | 9.455 | 9.067 | 10.019 | 10.555 | 9.816 | 10.422 | 10.428 |
| Pax6-Up | AK086933 | Fas | 5.428 | 6.446 | 7.171 | 5.344 | 5.492 | 5.385 | 5.389 | 5.537 | 5.416 | 6.349 | 6.558 | 6.362 | 6.748 | 6.211 | 7.364 | 7.017 | 7.419 | 7.048 | 7.005 |
| Pax6-Up | AK086789 | AK086789 | 9.995 | 10.496 | 11.134 | 10.113 | 10.207 | 9.894 | 9.695 | 9.965 | 10.098 | 10.545 | 10.546 | 10.260 | 10.515 | 10.613 | 10.761 | 11.199 | 11.807 | 10.860 | 11.044 |
| Pax6-Up | AK085847 | AK085847 | 5.579 | 7.223 | 7.851 | 5.656 | 5.671 | 5.607 | 5.580 | 5.510 | 5.451 | 7.089 | 7.267 | 7.184 | 7.273 | 7.300 | 8.042 | 7.655 | 7.913 | 8.062 | 7.584 |
| Pax6-Up | NM_001079883 | Bcl11b | 7.716 | 8.455 | 10.554 | 7.837 | 8.006 | 7.575 | 7.790 | 7.324 | 7.765 | 8.352 | 8.524 | 8.504 | 8.696 | 8.200 | 10.991 | 10.693 | 10.270 | 10.237 | 10.578 |
| Pax6-Up | NM_023158 | Cxcl16 | 8.647 | 9.194 | 9.857 | 8.498 | 8.781 | 8.657 | 8.708 | 8.449 | 8.792 | 9.069 | 9.257 | 9.504 | 9.135 | 9.005 | 10.453 | 9.513 | 9.779 | 9.987 | 9.555 |
