## Supplementary Table-3 for "Loss of Corneal Raver2 Expression in Aniridia Associated Keratopathy"

**Supplemental Table 3. Patient data corresponding to Figure 2**

| Normal/Diseased Cornea # | Phenotype | Identifier | Age | Gender | Source |
|---|---|---|---|---|---|
| 1 | Normal | S08-1117 | 50 | F | UCL A |
| 2 | Normal | OP-12-21 | 91 | F | University of Virginia |
| 3 | Normal | 12-1088-10 | 17 | M | Utah Lions Eye Bank |
| 4 | Normal | 13-0468-100 | 17 | F | Utah Lions Eye Bank |
| 5 | Normal | 13-1023-100 | 22 | F | Utah Lions Eye Bank |
| 6 | Normal | 13-0167-205 | 39 | M | Utah Lions Eye Bank |
| 1 | Aniridia | 101347 | 36 | F | University of Virginia |
| 2 | Aniridia | OP12-188 | 39 | M | Indiana University |
| 3 | Aniridia | OP13-156 | 41 | M | Indiana University |
| 4 | Aniridia | 04-1004 | 26 | M | KKESH, Saudi Ara bia |
| 5 | Aniridia | 894 | 32 | F | University of Uta h |
| 6 | Aniridia | OP11-1901 | 41 | M | University of Virginia |
| 7 | Aniridia | S10-16146 | 17 | M | UCLA |
| 8 | Aniridia | 81219 | 32 | F | University of Virginia |
| 9 | Aniridia | 77739 | 42 | F | Wills Eye Inst itute |
| 10 | Aniridia | S09-01594 | 46 | F | UCLA |
